# NucleiSky enables cross-scale multimodal registration of microscopy data using nuclei constellations

**DOI:** 10.64898/2026.06.29.735028

**Authors:** Iván Hidalgo Cenalmor, Adán Olguín-Olguín, Carolina Prieto, Johannes Kumra Ahnlide, Pontus Nordenfelt, Ricardo Henriques, Mario Del Rosario, Guillaume Jacquemet

## Abstract

Integrating tissue-level organisation with sub-cellular resolution and molecular information often requires combining multiple microscopy modalities and scales. However, aligning images acquired with different modalities, settings, or instruments remains challenging. Here, we introduce NucleiSky, a microscopy image registration framework that utilises the spatial arrangement of nuclei or other landmarks as an intrinsic biological fingerprint. NucleiSky represents images as constellations of centroids and aligns them using geometric algorithms and spatial consensus scoring. In benchmark datasets, NucleiSky could localise query regions within larger reference images using as few as five nuclei. We show that NucleiSky can locate high-magnification fields of view within low-magnification overview scans, map these alignments to additional channels, support live brightfield-to-fixed registration using synthetic nuclear labels, and guide microscope retargeting. We further show that the same constellation-matching principle can be extended to 3D localisation and to non-nuclear landmarks. These findings establish local landmark geometry as an intrinsic spatial fingerprint that enables localisation and registration across imaging scales, modalities and microscopy platforms. NucleiSky is available as an open-source Python package and as notebook-based applications.

## Introduction

Biological imaging increasingly relies on integrating information across modalities and scales. In correlative and multi-modal imaging workflows, a reference image is often acquired first, for example, from a tissue section, stitched mosaic or stack, and then used to guide more specialised measurements. These may include high-resolution imaging using confocal microscopy, super-resolution microscopy, electron microscopy, or atomic force microscopy, as well as higher-content molecular readouts, including multiplexed protein imaging and spatial omics approaches (Enninful et al., 2026; Goltsev et al., 2018; Kim et al., 2023; Krentzel et al., 2025; Semba and Ishimoto, 2024). A related re-localisation problem arises when the same field of view must be revisited multiple times. For example, live-to-fixed imaging can link dynamic behaviours to endpoint molecular states (Almada et al., 2019), while iterative staining enables multiplexed imaging by repeatedly imaging the same field of view after sequential rounds of labelling (Lin et al., 2015; Radtke et al., 2020). Across these workflows, a central requirement is to align images acquired under different conditions, so that molecular, mechanical, or ultrastructural measurements can be interpreted together.

Although the image-registration field is mature (Fernandez and Moisy, 2021; Guizar-Sicairos et al., 2008; Klein et al., 2010; Pylvänäinen et al., 2023; Saraiva et al., 2025a; Thevenaz et al., 1998; Yaniv et al., 2018), aligning microscopy images across different scales and modalities without external fiducials can remain a significant challenge. Intensity-based registration frequently fails when appearances differ between modalities, such as hematoxylin and eosin (H&E) versus fluorescence, or electron microscopy versus light microscopy. Feature-based methods are vulnerable to staining differences, section distortions, and the complexity of searching extensive tissue areas. Consequently, multimodal registration often relies on manual landmark selection, grid dishes, and repeated trial and error, which reduce throughput, reproducibility, and the feasibility of automated microscopy workflows.

Cell organelles provide valuable opportunities for aligning microscopy images. For instance, mitochondria have proven useful for aligning high-resolution fluorescent images with FIB-SEM datasets (Krentzel et al., 2025). Interestingly, nuclei can serve as widely accessible intrinsic fiducials. They are common in biological samples, show distinct local arrangements, and can be easily stained with dyes or inferred via label transfer (Follain et al., 2026; von Chamier et al., 2021). For example, nuclei have been successfully used to align serial histopathology slices (Jeyasangar et al., 2024; Nasir et al., 2025). Additionally, nuclei are typically straightforward to segment using conventional image-processing techniques or deep-learning workflows (Hollandi et al., 2020; Pachitariu et al., 2025; Schmidt et al., 2018).

Inspired by astronomical methods that identify unknown images by matching star asterisms despite unknown translation, rotation, and scale, we developed NucleiSky to use nuclei as a modality-agnostic coordinate system for microscopy registration (Beroiz et al., 2020; Groth, 1986; Lang et al., 2010; Pál and Bakos, 2006). NucleiSky performs localisation and registration by matching constellations of nuclei (or other landmarks) between a query image, such as a high-resolution crop, and a larger reference image, such as a low-resolution overview or tiled scan. Recognising that no single approach is likely to capture local landmark geometry across all experimental settings, NucleiSky incorporates multiple complementary constellation matchers that represent neighbourhood structures in distinct ways. These matchers propose candidate similarity transforms based on landmark positions or, when available, per-landmark features, and evaluate them using a spatial-consensus step. This design enables robust matching across sparse constellations, repetitive patterns, and large search spaces, allowing for the localisation of queries with as few as five nuclei under favourable conditions. To accommodate diverse datasets without manual parameter tuning, NucleiSky also provides an adaptive matching strategy that prioritises rapid initial searches and increases computational effort only when necessary, thereby supporting practical cross-scale navigation across large microscopy datasets.

## Results

### The NucleiSky pipeline

In astronomy, an unknown field of view can be mapped onto a star chart by matching the relative arrangement of stars. Analogously, nuclei can serve as biological landmarks for aligning microscopy images (Jeyasangar et al., 2024). Nuclei are prevalent across tissues and samples, can be visualised using various methods, and are generally straightforward to segment. NucleiSky applies this constellation-based approach to locate a query field of view within a larger reference image and to estimate the similarity transform that aligns the two sets of nuclear points (Fig. 1).

**Fig. 1.**
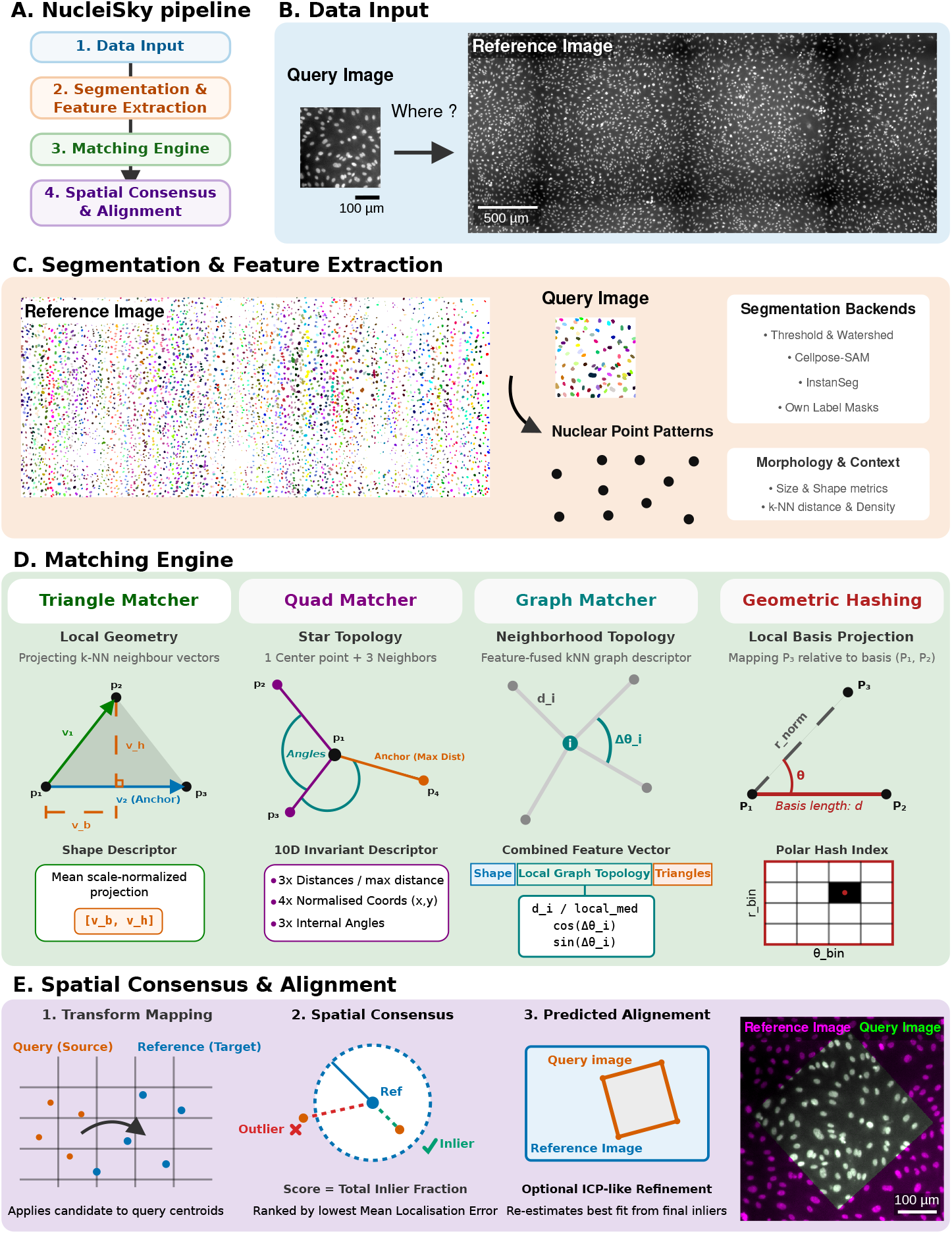
Overview of the NucleiSky processing pipeline. (**A**) Overview of the NucleiSky workflow. (**B**) A query field of view (or crop) is localised within a larger reference image. The query can be acquired at a different scale and in a different modality from the reference. (**C**) Nuclei are segmented using user-supplied masks, a classical threshold-and-watershed pipeline, or deep learning-based backends (such as Cellpose-SAM or InstanSeg). The segmentation is reduced to a constellation of nuclear centroids and optional per-nucleus descriptors that serve as a modality-agnostic coordinate system. Centroids provide geometry, while morphology and neighbourhood features help disambiguate repetitive nuclear patterns. (**D**) Matching engine. Multiple complementary matchers propose candidate similarity transformations based on local nuclear configurations. These matchers trade off robustness and speed across regimes of landmark density, segmentation noise and search-space size, using graph-based matching, triangle- and quad-based geometric descriptors, or geometric hashing. In the schematics, *p* denotes landmark points, and *v* denotes vectors. For the triangle matcher, *v*_*b* and *v*_*h* denote the normalised base-projection and signed-height components of the local triangle descriptor. For the graph matcher, *d*_*i* denotes the distance from a nucleus to its ith local neighbour, and Δ*θ*_*i* denotes the relative angular offset of that neighbour; distances are normalised by the median local neighbour distance, local_med. For geometric hashing, r_norm denotes the normalised distance of *p*_3_ in the local coordinate frame, and *θ* denotes its angular coordinate. (**E**) Candidate transforms are then ranked by spatial consensus after mapping the query constellation into the reference frame. Matched nuclei form an inlier set within a fixed tolerance, enabling the selection and optional refinement of the best transform. NucleiSky returns the final alignment alongside the predicted bounding box for visual validation and downstream analysis. Each candidate registration is accompanied by quantitative confidence metrics, including the inlier fraction and localisation error. This approach enables users to distinguish reliable registrations from ambiguous matches.

The NucleiSky workflow (Fig. 1A) begins by assembling landmark constellations in both the query field and the larger reference image, either by detecting nuclei from existing label masks or by segmenting them using one of the built-in segmentation backends (Fig. 1B and C). Since nuclear segmentation is a common image analysis task, many methods exist, including several deep learning models that perform well across diverse images. We implemented a classical threshold-and-watershed workflow in the NucleiSky segmentation backends, alongside popular deep learning models such as Cellpose-SAM (Pachitariu et al., 2025) and InstanSeg (Goldsborough et al., 2024). When segmentation occurs within NucleiSky, and the nuclear size varies widely across inputs, images can optionally be rescaled before detection to normalise nuclear size, thereby enhancing robustness for scale-sensitive segmentation models (Ferreira et al., 2025). From the resulting masks, NucleiSky extracts nuclear centroids and, when possible, per-nucleus descriptors that summarise morphology and neighbourhood context (for example, size and shape descriptors and local neighbour statistics). These descriptors are then standardised and combined with rotation and scale-normalised geometric encodings used by the matchers.

Rather than comparing image intensities directly, NucleiSky reduces each image to a geometric representation whose identity is preserved across imaging modalities. The query constellation must be aligned with the reference coordinate system despite unknown translations, rotations, and scales. We implemented four matchers in NucleiSky. These matchers differ in how they generate candidate similarity transforms from local geometric evidence (Fig. 1D). The graph matcher fuses per-nucleus feature vectors with graph- and triangle-derived local geometry; the triangle matcher uses local triangles for matching; the quad matcher uses four-point configurations to produce rotation- and scale-invariant descriptors; and the geometric hashing matcher builds an index of local reference frames. Each matcher suggests plausible correspondences, converts them into transform hypotheses, and then uses the same consensus strategy to evaluate candidates. After applying a candidate transform to the query centroids, NucleiSky identifies inliers as transformed nuclei that fall within a fixed tolerance of a reference nucleus. It then scores hypotheses based on spatial inlier support and reports the inlier fraction and mean localisation error (Fig. 1E). The best transform can optionally be refined via iterative updates. The final output includes both the estimated alignment and the predicted query footprint, represented as a bounding box in reference pixel coordinates, enabling quick verification and downstream analysis (Fig. 1E).

### Benchmarking of the NucleiSky pipeline

With the NucleiSky pipeline in place, we next asked whether landmark constellations contain sufficient spatial information to localise a small query field of view within a larger reference image. To test this, we cropped query patches of increasing size from four datasets and used NucleiSky to recover the position of each patch in the original image (Fig. 2; Methods). Datasets 1 and 2 comprised tiled images of fixed DAPI-stained endothelial monolayers, with nuclei segmented either by classical intensity thresholding (Dataset 1; 4,387 nuclei; 1,266 × 2,970 pixels) or by Cellpose-SAM (Dataset 2; 14,418 nuclei; 5,520 × 5,520 pixels). Dataset 3 was a substantially larger bright-field whole-slide image of an H&E-stained mouse colon section, segmented with InstanSeg (247,107 nuclei; 32,201 × 28,329 pixels), providing a more challenging search space. Dataset 4 tested whether the same principle could extend beyond nuclei by using a tiled image of DAPI-stained E. coli containing 81,379 detected bacteria, segmented with Cellpose-SAM.

**Fig. 2.**
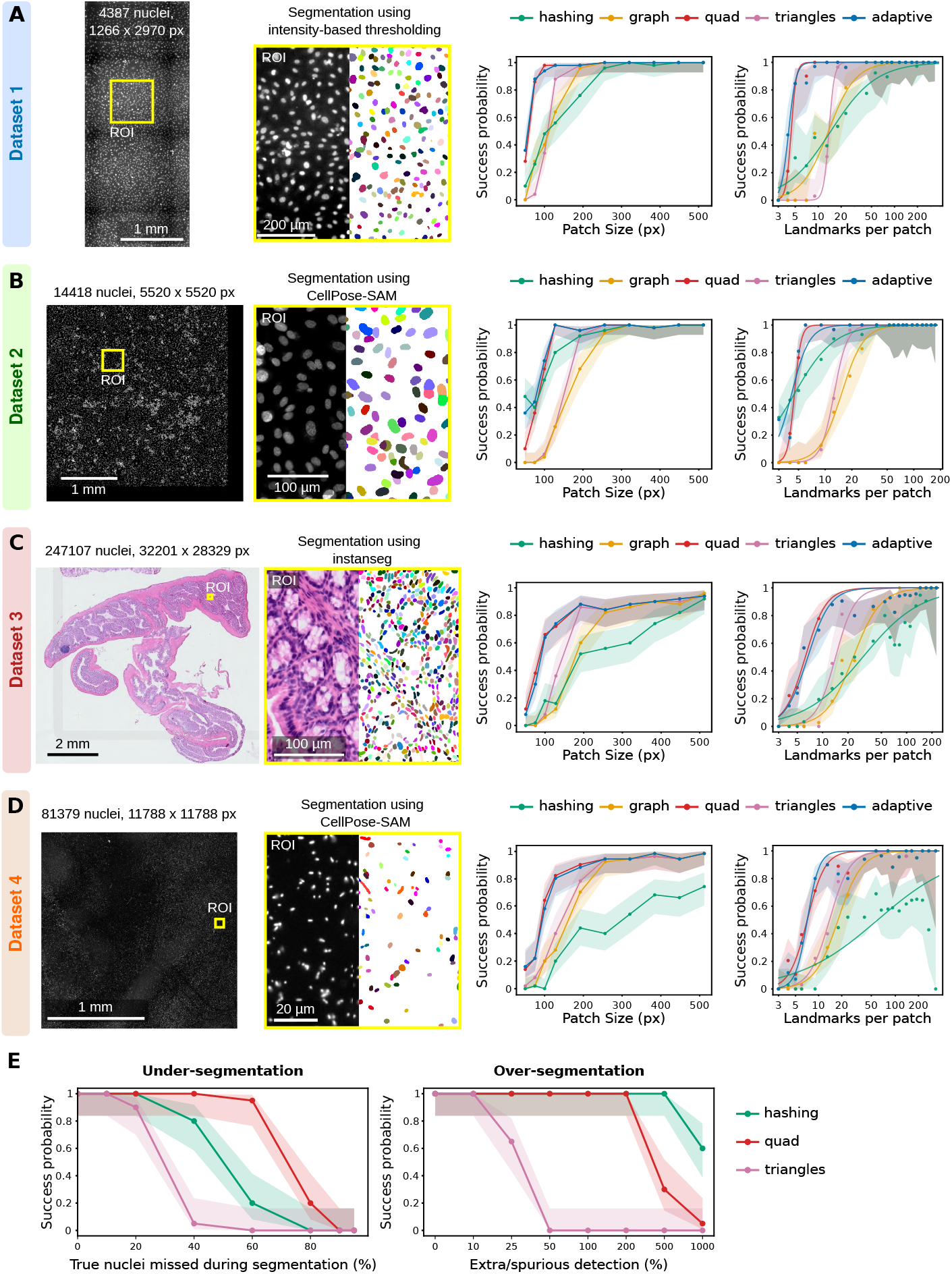
Benchmarking NucleiSky localisation performance. (**A**–**D**) Evaluation of NucleiSky’s ability to localise small query fields of view within larger reference images using landmark constellations. Query patches ranging from 50 to 512 pixels were cropped from each reference image and then re-localised with NucleiSky. For each dataset, the panels show, from left to right: the full reference image, a magnified region of interest, the corresponding segmentation mask, the localisation probability as a function of query patch size, and the localisation probability as a function of the number of detected landmarks in the query field of view. (**A**) Dataset 1: tiled image of a fixed DAPI-stained endothelial monolayer, measuring 1,266 × 2,970 pixels and containing 4,387 nuclei segmented using classical intensity thresholding. (**B**) Dataset 2: larger tiled image of a DAPI-stained endothelial monolayer, measuring 5,520 × 5,520 pixels and containing 14,418 nuclei segmented using the Cellpose-SAM backend. (**C**) Dataset 3: bright-field whole-slide image of an H&E-stained mouse colon section, measuring 32,201 × 28,329 pixels and containing 247,107 nuclei segmented using InstanSeg. (**D**) Dataset 4: tiled image of DAPI-stained E. coli, measuring 11,788 × 11,788 pixels and containing 81,379 detected objects segmented with Cellpose-SAM. In the localisation performance plots, points represent the observed localisation probability, and shaded bands represent 95% Wilson confidence intervals. For these plots, successful localisation was defined as matcher-reported success, an inlier fraction ≥ 0.80, SSIM ≥ 0.80, and a translation error <20 pixels. Patch-size plots show success as a function of query field size in pixels, whereas landmark-count plots show success as a function of the number of detected landmarks in the query field of view. The landmark-count x-axis is shown on a logarithmic scale to highlight the low-landmark regime, where localisation limits are most apparent. Smooth curves indicate logistic fits in log10(landmark count) space and are shown only as visual summaries. (**E**) Segmentation-error robustness benchmark using Dataset 1. Nuclei centroid sets were systematically degraded by removing true nuclei to mimic missed detections or by adding synthetic centroids to mimic spurious detections. Perturbations were applied to both the query and reference segmentations. The benchmark was conducted on 512×512-pixel query patches using the quad, triangle, and hashing matchers. Points indicate the observed probability of correct localisation, and shaded bands indicate 95% Wilson confidence intervals. For this benchmark, success was defined solely by image-level agreement between the original query crop and the predicted reference crop (SSIM ≥ 0.80).

Across datasets, localisation depended primarily on the uniqueness of the local landmark geometry rather than on image appearance or modality. Once a sufficient number of landmarks were present, however, localisation became highly reliable (Fig. 2). To compare matchers, we summarised performance using the fitted landmark threshold required to reach 90% localisation probability (Table 1). In Dataset 1, the quad matcher reached this threshold with as few as 5 nuclei, whereas the triangle, graph, and hashing matchers required 16, 26, and 44 nuclei, respectively. Dataset 2 showed a similar pattern: quad reached 90% success with approximately 5 nuclei, while triangle and graph required 17.5 and 25.1 nuclei, respectively; in this dataset, hashing reached the same threshold with 11.4 nuclei. In the larger H&E whole-slide image, all matchers required larger constellations, consistent with the increased search space and more repetitive tissue architecture. Nevertheless, quad and triangle remained the most sample-efficient, achieving 90% success with 9.6 and 18.3 nuclei, respectively, whereas graph and hashing required approximately 40 and 94 nuclei, respectively. In the E. coli dataset, quad again required the fewest landmarks (12.7 objects), followed by triangle (27 objects) and graph (38.8 objects), while hashing did not reach the 90% success threshold.

**Table 1.** Estimated landmark thresholds for 90% localization probability across matchers and benchmark datasets. Each value reports the fitted number of landmarks required to achieve a 90 percent probability of localisation. Thresholds were estimated from logistic fits in log_10_(landmark count) space, as used for the landmark-count plots in Fig. 2. Hashing did not reach the 90% localization threshold in Dataset 4 and is reported as n/a.

| Dataset | Triangle | Quad | Graph | Hashing |
| --- | --- | --- | --- | --- |
| Dataset 1 | 16 | 5 | 26 | 44 |
| Dataset 2 | 17.5 | 5 | 25.1 | 11.4 |
| Dataset 3 | 18.3 | 9.6 | 40 | 94 |
| Dataset 4 | 27 | 12.7 | 38.8 | n/a |

We next assessed runtime under the same benchmark conditions. Localisation was generally fast on a standard laptop workstation, with search times below 1 s for Dataset 1 and approximately 2 s for Dataset 2. By contrast, the much larger H&E reference required approximately 20 s per query (Fig. S1A; Methods). Runtime also varied by matcher, with the triangle matcher generally providing the fastest searches (Fig. S1A). These results indicate that constellation-based localisation is computationally practical for tiled microscopy images and remains feasible even with whole-slide-scale references.

Because segmentation quality can vary, we next tested NucleiSky’s sensitivity to segmentation errors. Using Dataset 1, we selected 512 × 512-pixel query patches containing at least 50 nuclei and degraded both the query and reference centroid sets. To do so, we removed true nuclei and added synthetic centroids to mimic missed or spurious detections, respectively (Fig. 2E; Methods). We evaluated the quad, triangle, and hashing matchers individually and defined success solely by image-level recovery of the correct field of view, measured by SSIM. Under this stringent condition, the quad matcher was most robust to missing nuclei, maintaining at least 90% success even with up to 60% landmark loss, compared with 20% for both the hashing and triangle matchers. Conversely, hashing was most tolerant of spurious detections, maintaining at least 90% success up to 500% extra centroids, compared with 200% for quad and 10% for triangles.

Because benchmarking indicated that no single matcher was optimal across all datasets, we implemented an adaptive controller to reduce the need for manual matcher selection (Fig. 2). The adaptive mode selects and escalates matchers based on the number of query landmarks and the availability of feature vectors, prioritising matchers suited to small constellations and escalating when additional search effort is required (Methods).

Collectively, these results demonstrate that relatively small groups of nuclei contain sufficient geometric information to uniquely identify their spatial context within much larger images, even across datasets that differ substantially in imaging modality, landmark density, and scale.

### Localising high-magnification fluorescence fields of view within low-magnification tile scans

Having established NucleiSky’s baseline localisation performance under controlled benchmarking conditions, we next tested whether the same approach could support practical microscopy re-targeting scenarios. In these settings, query and reference images are often acquired under different conditions, including different magnifications and stage orientations. We therefore asked whether NucleiSky could localise high-magnification fluorescence fields of view within low-magnification overview tile scans. To do so, we used data previously prepared for a correlative light- and electron-microscopy workflow on confluent epithelial monolayers of DCIS.com cells (Grobe et al., 2026; Miller et al., 2000; Peuhu et al., 2022) (Fig. 3).

**Fig. 3.**
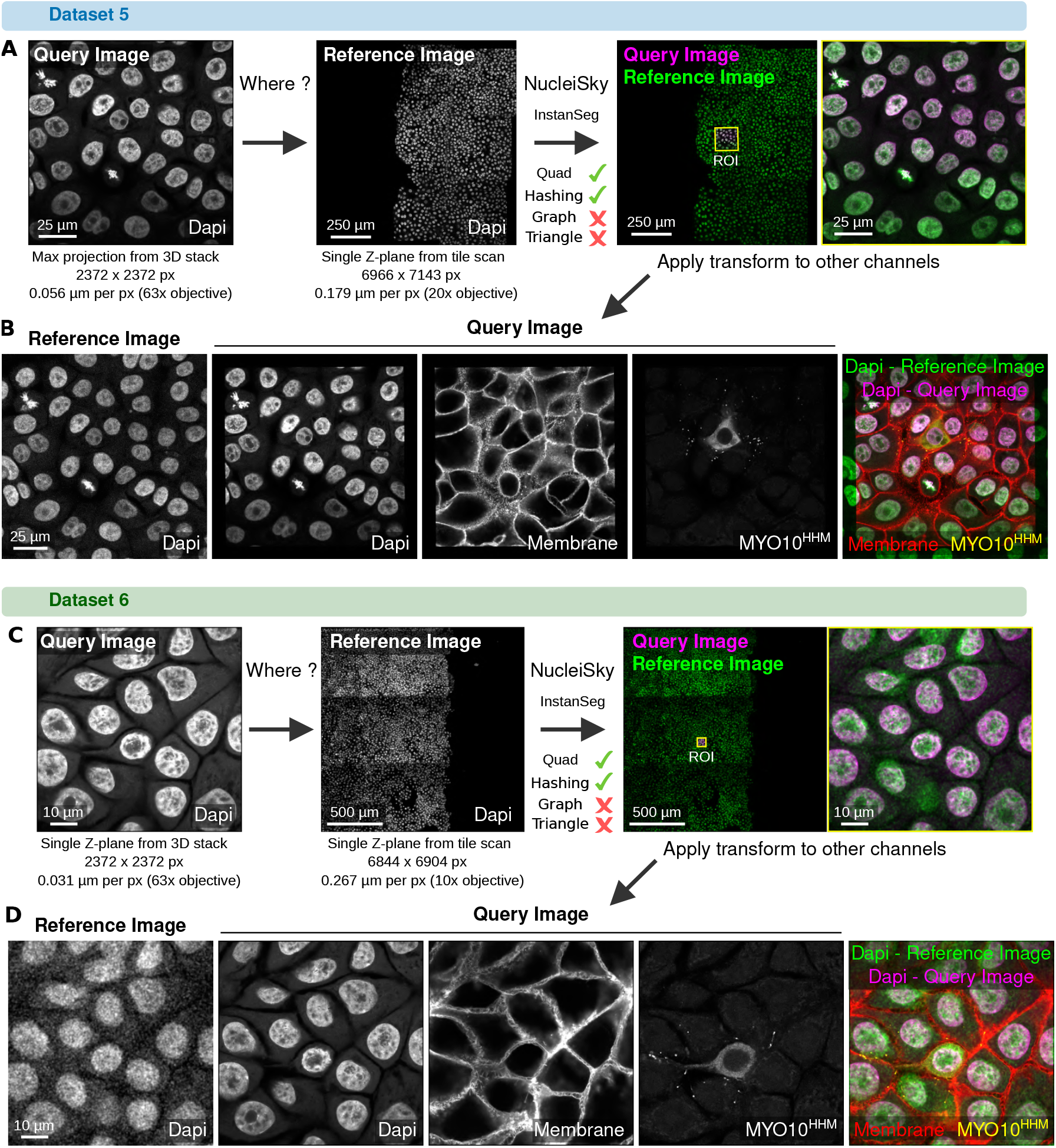
Cross-scale localisation of high-magnification acquisitions within overview tiles. (**A**) Dataset 5. A high-magnification query (maximum-intensity projection from a 63× stack) is localised within a lower-magnification 20× tile-scan reference acquired from a confluent DCIS.com epithelial monolayer. From left to right: the query and reference inputs, the predicted query footprint overlaid on the full reference, and a magnified view of the matched region (reference in green, query in magenta). This highlights the consistent nuclear geometry despite differences in sampling and a slight relative rotation. The overlay is rendered in reference-pixel units, with the query downsampled to match the reference sampling. (**B**) Dataset 5, channel transfer. The saved transform from (A) is applied to additional query channels to position high-magnification molecular readouts within the reference context. From left to right: reference ROI DAPI, registered query DAPI, registered membrane channel, registered MYO10^HMM^-GFP signal (a truncated Myosin-X construct used as a filopodia-tip marker), and a merged view. The panel is displayed in reference-pixel units (query downsampled), with 20-pixel padding around the query footprint to visualise the monolayer’s continuity beyond the original crop. (**C**) Dataset 6. A higher-magnification 63× query is localised within a 10× tile-scan reference, demonstrating successful placement even when the reference contains less detail per unit area. As in (**A**), panels show the inputs, the predicted footprint on the full reference, and a magnified overlay of the matched region (reference in green, query in magenta). Here, the overlay is rendered in query-pixel units, with the reference crop upsampled to match the query’s sampling. (**D**) Dataset 6, channel transfer. The transform estimated in (**C**) is reused to register additional query channels onto the matched reference ROI. Images are displayed in query-pixel units, with the reference crop upsampled. All images were acquired on an Airyscan confocal microscope in standard super-resolution mode.

In the first example (Dataset 5), the query was a maximum-intensity projection from a high-magnification 3D stack acquired with a 63× objective (2,372 × 2,372 pixels; 0.056 µm/pixel). The reference was a single Z-plane from a 20× tile scan (6,966 × 7,143 pixels; 0.179 µm/pixel) (Fig. 3A, B). In the second example (Dataset 6), the query was captured at an even higher magnification from a 63× stack (2,372 × 2,372 pixels; 0.031 µm/pixel), whereas the reference was a lower-magnification 10× tile scan (6,844 × 6,904 pixels; 0.267 µm/pixel) that provided fewer structural details per unit area (Fig. 3C, D). In both datasets, nuclei were segmented using InstanSeg, yielding comparable centroid representations despite the drastic differences in sampling resolution.

Across these cross-scale comparisons, the quad and geometric hashing matchers produced accurate transformations, whereas the graph and triangle matchers failed to do so (using default settings). Notably, the query regions in both datasets showed a slight rotation relative to the reference, reflecting common experimental conditions. Once the transformation was estimated using the nuclear constellation, the same mapping was applied to other channels from the corresponding acquisitions. This enabled multimodal overlays, rendered either in the coordinate system and pixel size of the reference image (Fig. 3B) or in the query image’s coordinate system (Fig. 3D).

Collectively, these examples illustrate that NucleiSky can localise high-resolution fluorescence fields of view within low-resolution overview scans.

### Live-to-fixed registration using synthetic nuclear labels

We next tested NucleiSky in a second re-localisation scenario, aligning live-cell imaging video with endpoint stainings obtained after fixation. To assess whether NucleiSky could support a live-to-fixed imaging workflow, we used bright-field movies of cancer cells attaching to endothelial cells under controlled flow conditions in microfluidic devices (Follain et al., 2026). In these experiments, live imaging was performed in bright-field to avoid fluorescence phototoxicity. After live acquisition, samples were fixed and stained using a cell painting kit, including a nuclear channel, and imaged as fluorescence tile scans (Fig. 4A).

**Fig. 4.**
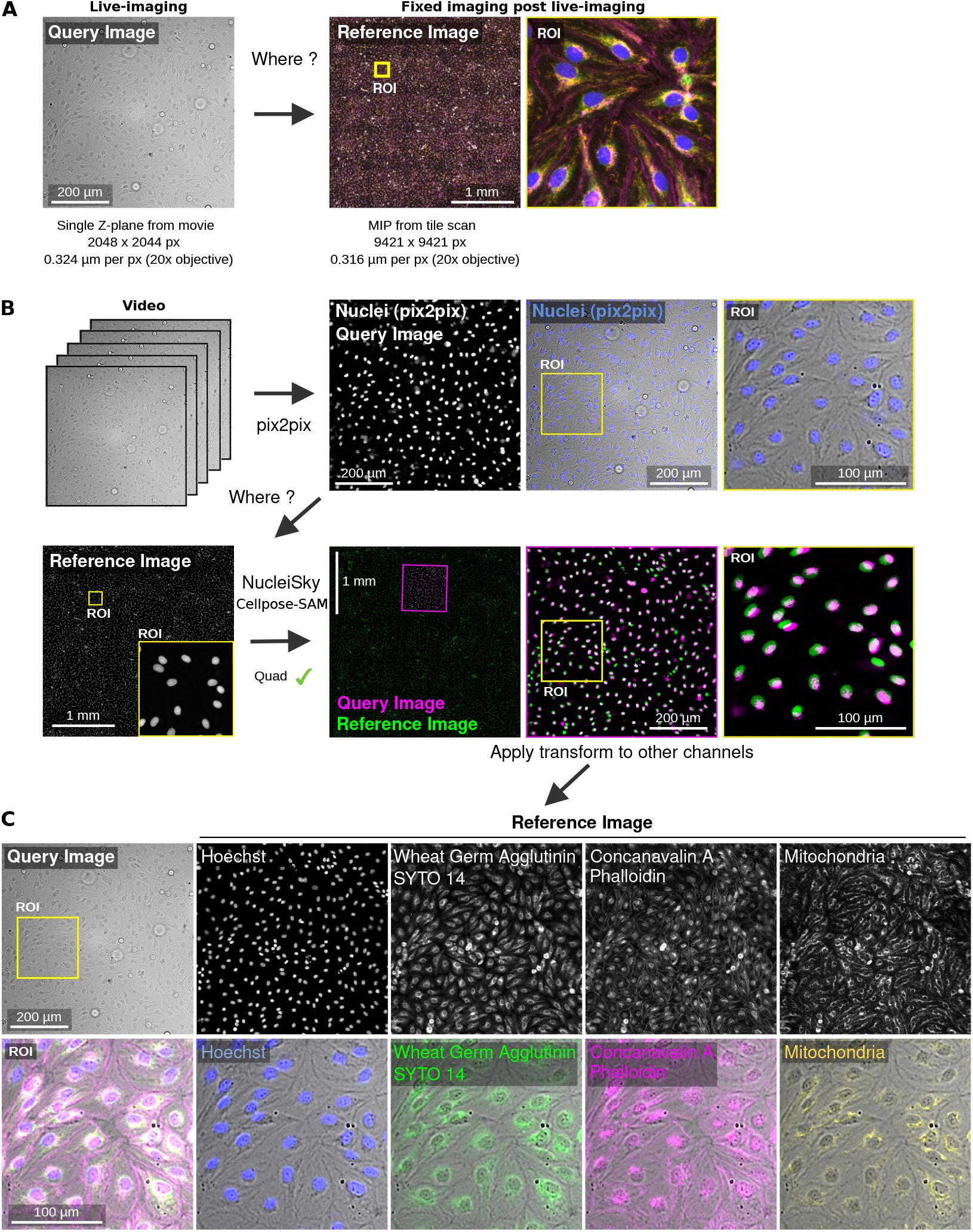
Live-to-fixed registration using bright-field-derived nuclear constellations. (**A**) Input images for live-to-fixed registration. The live query was taken from the final frame of a transmitted-light bright-field movie acquired during cancer-cell perfusion over an HPMEC monolayer in a microfluidic channel. After live imaging, the same sample was fixed, stained using a Cell Painting protocol, and re-imaged by spinning-disk confocal microscopy. The fixed reference was generated as a maximum-intensity projection. The query measured 2,048 × 2,044 pixels at 0.3245 µm per pixel, and the reference measured 9,421 × 9,421 pixels at 0.3169 µm per pixel. (**B**) Synthetic nuclear landmark generation and NucleiSky matching. A pix2pix artificial-labelling model was used to infer a DAPI-like nuclear image from the live bright-field input. The pix2pix-derived query nuclei and Hoechst-positive reference nuclei were segmented using the Cellpose-SAM backend in NucleiSky. The query constellation was matched to the fixed reference constellation using the quad matcher. Yellow squares highlight regions of interest, which are magnified. The magenta square highlights the query image within the reference image. (**C**) Application of the live-to-fixed transform to endpoint fluorescence channels. The similarity transform estimated from the nuclear constellations was applied to fixed-sample fluorescence channels (as indicated) acquired after live imaging. The yellow square highlights a region of interest, which is magnified.

Because no fluorescent nuclear channel was acquired during live imaging, we generated synthetic nuclear landmarks from the bright-field movie using a pix2pix artificial-labelling model developed for FlowVision (Follain et al., 2026; Isola et al., 2016). The model inferred a DAPI-like nuclear image from the transmitted-light input, producing a synthetic nuclear channel. The pix2pix-derived nuclei image and the fixed Hoechst reference were both segmented with Cellpose-SAM to generate the query and reference constellations. NucleiSky then matched these two constellations and estimated a similarity transform that maps the live field of view into the fixed fluorescence coordinate system (Fig. 4B). In this example, the quad matcher successfully localised the live-imaged region within the fixed reference tile scan.

Finally, the same transform was applied to additional endpoint fluorescence channels acquired after fixation (Fig. 4C). This linked the live bright-field field of view to post-fixation Cell Painting stainings, including membrane, cytoskeletal and mitochondrial labels. Thus, using synthetic nuclear labels inferred from label-free live imaging, NucleiSky can determine where the live imaging was performed in the sample, recover the corresponding fixed-cell field of view across changes in both modality and microscope system, and transfer the alignment to multiplexed endpoint molecular readouts. This approach demonstrates that NucleiSky can operate even when nuclei are never experimentally labelled during live imaging, provided that biologically meaningful landmarks can be computationally inferred. This provides a strategy to correlate dynamic behaviours observed during live imaging, such as cancer-cell arrest under flow, with cellular and molecular organisation acquired post-fixation.

### Smart microscopy using nuclei constellations

After demonstrating that NucleiSky can localise microscopy fields of view post-acquisition, we next tested if the same strategy could be applied directly on the microscope to guide acquisition. We integrated NucleiSky into a Nikon NIS-Elements JOBS workflow, in which NIS-Elements controls stage movement, autofocus, and image acquisition, while NucleiSky performs target recognition based on nuclear constellations (Fig. 5; Video 1; see Methods).

**Fig. 5.**
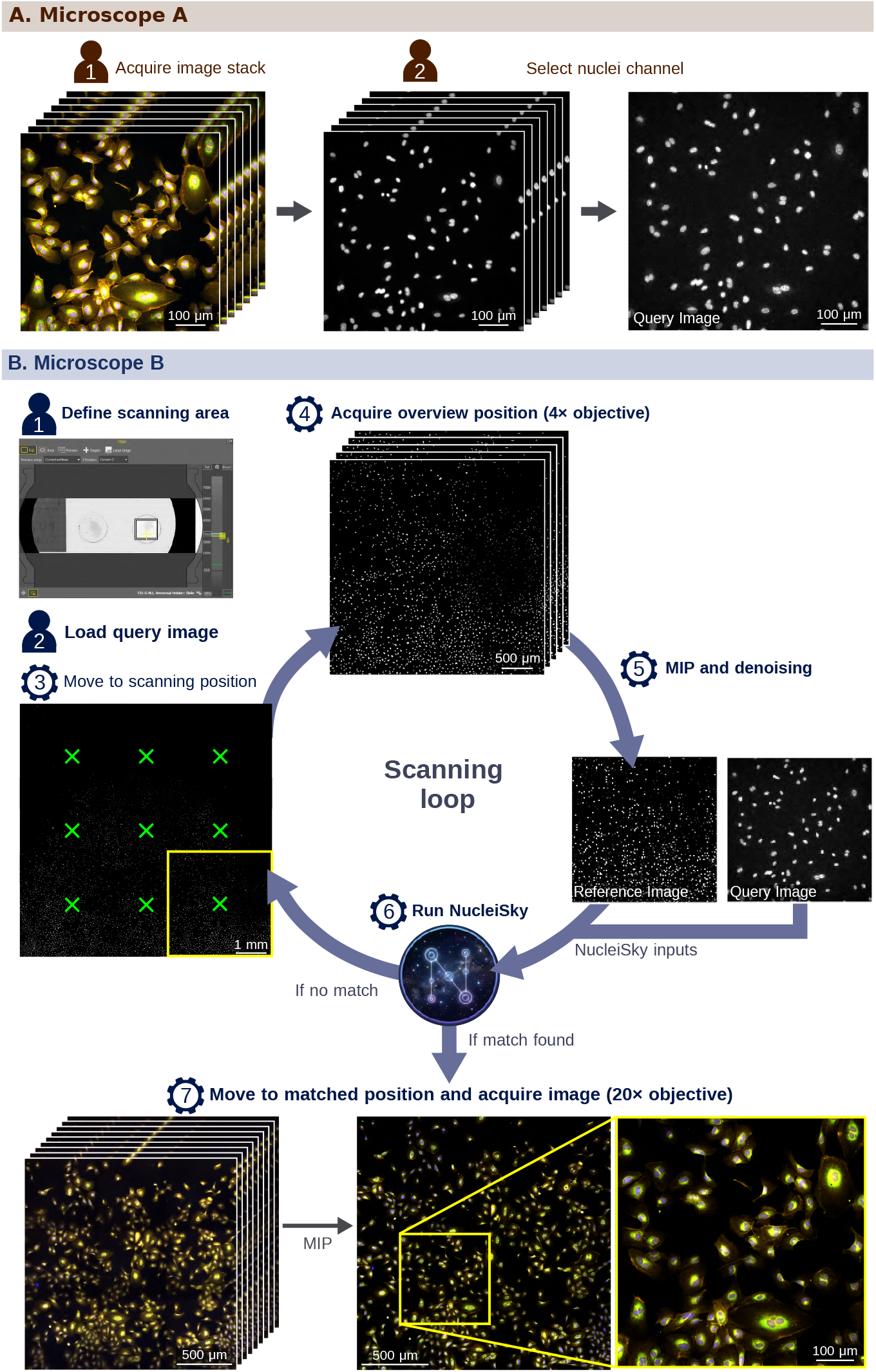
Smart microscopy using nuclei constellations to re-target fields of view across microscopes. (**A**) Query generation on Microscope A. A field of view is acquired on the first microscope, and the nuclei channel is selected from the image stack to generate the query image for downstream re-targeting. This query image contains the nuclear constellation that NucleiSky searches for on the second microscope. (**B**) Automated re-targeting workflow on Microscope B. The user defines a scanning area, loads the query image, and moves the stage to the starting scan position. The microscope then enters a scanning loop, acquiring low-magnification overview images with a 4× objective. At each position, the image stack is converted to a maximum-intensity projection, denoised, and passed to NucleiSky with the query image. If no acceptable match is found, the workflow advances to the next scan position. When NucleiSky identifies the query constellation, the microscope moves to the matched coordinates and acquires a higher-resolution image with a 20× objective. The final panels show successful re-localisation of the original field of view, with the matched region highlighted in the low-magnification overview and confirmed by the corresponding high-magnification acquisition.

The workflow begins by acquiring a field of view containing nuclei on the first microscope to generate the query image (Fig. 5A). This query contains the nuclear constellation to be re-identified later. The sample is then transferred to a second microscope, where the user defines a scanning area and loads the query image into the JOBS workflow (Fig. 5B). The microscope enters a low-magnification scanning loop, in which, at each position, it autofocuses and acquires an overview image stack with a 4× objective, converts the nuclei channel to a maximum-intensity projection, denoises the image, and passes the resulting reference image and the query image to NucleiSky. If no acceptable match is found, the stage advances to the next scanning position. When a match is detected, the loop stops, the stage moves to the predicted coordinates, performs a double-pass autofocus, and acquires a higher-resolution multichannel image with a 20× objective.

As a proof of concept, we used this workflow to image an endothelial monolayer stained with a Cell Painting kit on two microscopes. A region of interest was first acquired on a widefield microscope using a 20× objective. The same sample was then transferred to a spinning-disk confocal microscope, where the NucleiSky JOBS workflow performed low-magnification exploratory scanning and automatically identified the original nuclear constellation. The matched position was then re-imaged at higher resolution, confirming recovery of the original field of view on the second microscope (Fig. 5B; Video 1).

These results demonstrate that NucleiSky can serve as a real-time re-targeting engine for smart microscopy. By using nuclear constellations as persistent spatial signatures, the workflow enables fields of view to be re-identified across instruments or imaging sessions without manual searching, gridded coordinates, or external fiducials. This provides a path towards closed-loop acquisition workflows in which overview imaging, target recognition, and high-resolution re-imaging are linked automatically.

### Extending nuclei-constellation matching to volumetric datasets

We next asked whether constellation-based localisation could be extended from 2D images to volumetric microscopy data (Fig. 6A). In NucleiSky3D, each volume is represented as a constellation of nuclear centroids in calibrated 3D physical coordinates. NucleiSky3D retains the core architecture of the 2D pipeline, in which local landmark constellations identify candidate transforms, which are then evaluated via spatial consensus. For volumetric point clouds, we implemented two geometric matchers (Fig. 6B). The pyramid matcher extends triangle-based matching to 3D by constructing four-point tetrahedra from local k-nearest-neighbour relationships. The 3D geometric hashing matcher defines local coordinate frames from anchor nuclei and indexes neighbouring landmarks in 3D hash bins. Compatible configurations generate candidate transforms, enabling the selected transform to localise the query subvolume within the reference volume (Fig. 6C). As in the 2D pipeline, NucleiSky3D includes an adaptive controller that selects the matcher order based on query size.

**Fig. 6.**
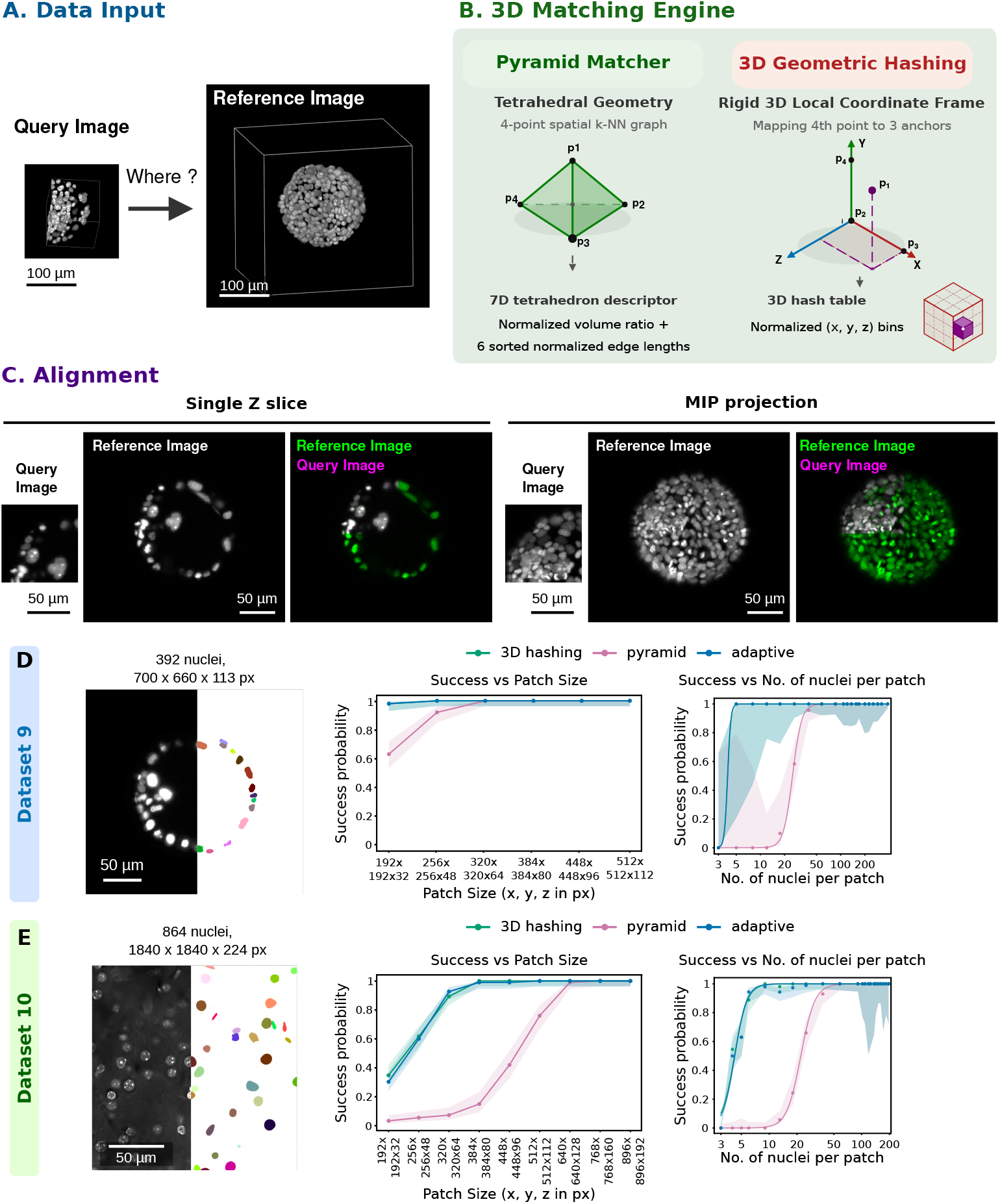
Extending nuclei-constellation matching to 3D stacks. (**A**) 3D data input. A query volume is localised within a larger reference volume by reducing both image stacks to calibrated 3D nuclear-centroid constellations. The example panels show the reference and query images, with scale bars indicated. (**B**) 3D matching engine. NucleiSky3D implements two volumetric geometric matchers. The tetrahedron-based pyramid matcher constructs local four-point configurations from a 3D k-nearest-neighbour graph and encodes each non-degenerate tetrahedron using a normalised volume term and six sorted, normalised edge lengths. The 3D geometric hashing matcher defines a local coordinate frame from three anchor nuclei and stores the normalised position of a fourth nucleus in a 3D hash table. This enables the retrieval of compatible local configurations and the generation of candidate 3D similarity transforms. (**C**) Alignment between the query image and the reference image. An example of successful alignment between a random 3D crop from the Platynereis-Nuclei-CBG dataset and the full image. Successful alignment is demonstrated both with a single Z-plane and with a maximum-intensity projection. (**D**-**E**) Evaluation of NucleiSky3D’s ability to localise query subvolumes within larger reference volumes using 3D nuclear constellations. For each dataset, query subvolumes of increasing size were cropped from the reference volume and re-localised with the pyramid, 3D geometric hashing, and adaptive 3D matchers. The panels show a single z-plane of the dataset, along with the corresponding segmentation examples and benchmark performance as functions of query subvolume size and the number of detected landmarks in the query volume. (**D**) Dataset 9: Platynereis-Nuclei-CBG reference volume, measuring 700 × 660 × 113 pixels (X, Y, Z) and containing 392 segmented nuclei. (**E**) Dataset 10: DAPI-stained mouse brain slice acquired as a spinning-disk confocal Z-stack, measuring 1,840 × 1,840 × 224 pixels (X, Y, Z) and containing 864 segmented nuclei. In the 3D localisation performance plots, points represent the observed localisation probability, and shaded bands represent 95% Wilson confidence intervals. Successful localisation was defined as matcher-reported success, an inlier fraction 0.50, and a translation error <20 pixels. Patch-size plots show success as a function of query subvolume size in pixels, whereas landmark-count plots show success as a function of the number of detected nuclei in the query subvolume. The landmark-count x-axis is shown on a logarithmic scale to highlight the low-landmark regime, where localisation limits are most apparent. Smooth curves indicate logistic fits in log10(landmark count) space.

NucleiSky3D can use existing 3D label masks or generate landmarks from raw volumes using a 2.5D segmentation workflow. In this mode, individual planes are segmented with the same 2D backends described above, and detections are linked across adjacent Z-slices using an intersection-over-union criterion to reconstruct 3D nuclear objects.

To benchmark NucleiSky3D, we used the same patchlocalisation strategy as in 2D, extended to query volumes. We tested two datasets with different landmark densities and imaging characteristics. Dataset 9 was the publicly available Platynereis-Nuclei-CBG dataset (Lalit et al., 2022). The accompanying nuclear instance masks allowed us to evaluate matching independently of segmentation quality. This reference measured 700 × 660 × 113 pixels (X, Y, Z) and contained 392 segmented nuclei (Fig. 6D). Dataset 10 was a DAPI-stained mouse brain slice acquired as a spinning-disk confocal Z-stack, measuring 1,840 × 1,840 × 224 pixels and containing 864 segmented nuclei (Fig. 6E). For each dataset, query subvolumes of increasing size were cropped from the reference and re-localised using the pyramid, 3D geometric hashing, and adaptive matchers.

As in the 2D benchmark, performance was primarily determined by the number of landmarks in the query. In the Platynereis dataset, 3D geometric hashing was the most sample-efficient, reaching 90% localisation probability with 5 nuclei. The pyramid matcher required larger constellations, reaching the same threshold at 34 nuclei (Fig. 6D). In the mouse brain dataset, 3D geometric hashing again performed best in the low-landmark regime, requiring 6 nuclei to reach 90% localisation probability, whereas pyramid matching required 33 nuclei (Fig. 6E). Thus, across both datasets, adaptive and 3D geometric hashing could localise sparse subvolumes from only a few nuclei, whereas pyramid matching became reliable only when larger 3D constellations were available.

Runtime remained compatible with practical volumetric localisation. In the Platynereis benchmark, median total query times were approximately 0.12 s for adaptive mode, 0.12 s for pyramid matching, and 1.19 s for 3D geometric hashing. In the mouse brain benchmark, adaptive searches ranged from approximately 0.3 to 7 s across the tested patch sizes, whereas 3D geometric hashing was more stable, at approximately 2.6-3.9 s per query.

Together, these results demonstrate that NucleiSky3D extends constellation-based localisation from 2D images to volumetric data. Although the tested 3D references remain modest compared with whole-organ or cleared-tissue volumes, they establish the feasibility of geometry-based 3D localisation and provide a foundation for scaling to larger volumetric datasets.

## Discussion

Here, we introduce NucleiSky, an open-source toolbox that uses landmark constellations as a coordinate system for microscopy image registration. By reducing images to calibrated landmark constellations, NucleiSky localises small query fields of view within much larger reference images, achieving successful localisation from as few as five nuclei under favourable conditions. We show that this strategy enables cross-scale registration of high-magnification fluorescence images within low-magnification overview scans, transfer of the resulting transforms to additional channels, live-to-fixed registration, automated field-of-view re-targeting on the microscope, and also works with volumetric data. Together, these findings indicate that nuclei can serve not only as objects for segmentation and quantification but also as endogenous spatial landmarks for preserving image identity across scales, modalities, instruments, and dimensions.

NucleiSky draws inspiration from blind astrometry, in which an unknown astronomical field is identified from local star geometry despite unknown translations, rotations, and scales. This concept has a long history, from triangle-invariant matching of two-dimensional coordinate lists to Delaunay-based triangle spaces and Astrometry.net’s hashing of four- and five-star asterisms, followed by statistical verification against a reference catalogue (Groth, 1986; Lang et al., 2010; Pál and Bakos, 2006). A related software tool to NucleiSky is astroalign, which uses triangle invariants for direct star alignment (Beroiz et al., 2020). NucleiSky follows the same central idea, that local point geometry can identify a field of view, but applies it to a different scientific field.

Constellation-based point-cloud matching also has an important precedent in microscopy. Preibisch and colleagues demonstrated that multiview SPIM datasets can be registered by detecting fluorescent beads and matching their constellations using geometric descriptors, accelerated by geometric hashing and filtered by RANSAC (Preibisch et al., 2010). However, their task was to align multiple overlapping views of the same SPIM volume using externally introduced fiducials. NucleiSky addresses a different localisation problem: identifying where a small query field of view or subvolume belongs within a much larger reference image, even when acquired using different magnification and modalities. NucleiSky builds on the same broader principle that local landmark geometry can identify corresponding fields, but applies it to endogenous biological landmarks rather than fluorescent beads. Unlike beads, nuclei are dense and often locally repetitive. We therefore implemented multiple complementary matchers, spatial-consensus scoring, and adaptive matcher selection in NucleiSky to support localisation across microscopy samples, scales, and modalities.

Nuclei have previously been used as landmarks to support the alignment of serial tissue sections (Jeyasangar et al., 2024; Nasir et al., 2025). Typically, previous approaches use nuclei for local refinement after the corresponding tissue region has been approximately identified. For instance, a previous workflow converts detected nuclei into point sets and combines rigid alignment, non-rigid point-set registration, and image warping to align serial tissue sections (Jeyasangar et al., 2024). Another whole-slide registration framework, CORE, uses nuclear centroids and shape information to register images after a separate coarse alignment stage (Nasir et al., 2025). These methods establish nuclei as useful registration landmarks, but they require many nuclei to work; for example, one nuclei-location workflow recommends tiles containing approximately 100 nuclei (Jeyasangar et al., 2024). NucleiSky addresses a different problem. Rather than using nuclei only to refine already corresponding fields, it treats small groups of nuclei as searchable constellations to identify where a query field belongs within a larger reference. NucleiSky can therefore operate in a partial-overlap regime without requiring individual nuclei to be uniquely identifiable or the target region to be known in advance.

Retargeting the same field of view is a common challenge in microscopy. Existing strategies address this by preserving or externalising spatial identity. In live-to-fixed imaging, the sample is often kept on the microscope while treatment, fixation, and staining are performed in situ using microfluidics (Almada et al., 2019). Cyclic imaging methods such as CycIF, CODEX, and 4i similarly preserve correspondence by repeatedly imaging the same specimen through sequential staining or readout cycles (Goltsev et al., 2018; Lin et al., 2015; Gut et al., 2018).

In correlative light and electron microscopy, retargeting often relies on gridded coverslips, laser-inscribed or micromoulded coordinate systems, or fiducial markers that remain visible across modalities and support high-precision correlation (Karreman et al., 2015; Kukulski et al., 2011; Schorb and Briggs, 2014). These strategies are effective because they make spatial identity recoverable by design, but they require that the coordinate frame, carrier, or engineered landmarks remain valid. By using nuclear constellations as endogenous landmarks, NucleiSky provides another localisation strategy directly encoded in the sample itself. This sample-intrinsic approach reduces dependence on external fiducials, dedicated marker channels, or carrier-based coordinate systems. Because the core NucleiSky matching functions can be called from Python, NucleiSky can also be incorporated into microscope-control environments that expose Python interfaces, providing a route towards automated field-of-view recognition and re-targeting.

The current implementation of NucleiSky has several limitations. It estimates global similarity transforms in calibrated physical coordinates and therefore assumes that the relative arrangement of landmarks is largely preserved across images. This makes it well-suited for localisation and retargeting. However, it does not support affine shear, anisotropic scaling, or local non-rigid deformations introduced by moving cells, swelling, or tissue distortion. NucleiSky also depends on the quality and uniqueness of the landmark constellation. Our benchmarking indicates that NucleiSky can tolerate missing nuclei and spurious detections to some extent, but these will reduce confidence or produce ambiguous matches. Finally, although NucleiSky has been extended to 3D and benchmarked on two volumetric datasets, this mode remains less extensively validated than the 2D workflow. The current 3D implementation estimates global similarity transforms, relies on 3D centroid constellations generated either from existing masks or 2.5D slice-linked segmentation, and has so far been tested on modest reference volumes rather than whole-organ or cleared-tissue datasets. Future work will focus on scalable OME-Zarr-based workflows, lazy loading and chunked export, parallel-assisted search, and hybrid pipelines that combine constellation-based localisation with downstream deformable refinement.

Altogether, we envision constellation-based localisations as a general strategy for bridging microscopy modalities, enabling biological datasets acquired across instruments, resolutions, and experimental conditions to be linked through intrinsic spatial organisation rather than shared image appearance.

## Methods

### NucleiSky implementation

NucleiSky is implemented as a Python package, with separate 2D and 3D pipelines accessible via nucleisky2d and nucleisky3d (https://github.com/CellMigrationLab/NucleiSky). The repository includes reproducible Jupyter notebooks that demonstrate complete 2D and 3D registration workflows. The same functionality is available through a programmatic API suitable for batch processing and integration into acquisition or analysis workflows. NucleiSky depends on standard scientific Python libraries for array computing and data management, including NumPy (Harris et al., 2020) and pandas. It uses SciPy for spatial queries and numerical routines (Virtanen et al., 2020), and scikit-image for image processing utilities and resampling (van der Walt et al., 2014). Several matchers use NetworkX-based kNN graph representations (Hagberg et al., 2008) and Numba (Lam et al., 2015). Microscopy file I/O and export are handled by tifffile, while plotting utilities use matplotlib (Hunter, 2007). Optional dependencies include segmentation backends (Cellpose and InstanSeg), Zarr I/O for large volumetric datasets (Moore and Kunis, 2023), and SimpleITK to enhance 3D feature extraction (Moore and Kunis, 2023).

Additionally, we provide Jupyter notebooks to run NucleiSky2D and NucleiSky3D, along with associated benchmarks. These notebooks can be run in Google Colab or installed locally using the NucleiSky App, which was created with LabConstrictor (Hidalgo-Cenalmor et al., 2026).

### Image calibration, nuclei extraction, and constellation features

NucleiSky performs matching in physical units to support cross-magnification registration and reduce sensitivity to pixel sampling. Users provide the pixel size (µm/px) for 2D images and the voxel size for 3D volumes, or NucleiSky retrieves them from metadata when available. Centroid coordinates follow (y, x) in 2D and (z, y, x) in 3D. The estimated similarity transforms map query coordinates to the reference coordinate system. When built-in segmentation backends are used, images can optionally be rescaled before segmentation to ensure nuclei have comparable apparent diameters in pixels. This rescaling stabilises segmentation, whereas matching and export operate on centroids expressed in micrometres.

Nuclei segmentation can be performed using a threshold-and-watershed method or via learning-based techniques such as Cellpose (with a pretrained configuration called CellPose-SAM (Pachitariu et al., 2025)) or InstanSeg (Goldsborough et al., 2024). Alternatively, users may provide precomputed 2D or 3D label masks. For volumetric data, NucleiSky also supports a 2.5D approach that segments each Z-slice independently and then stitches labels across slices using intersection-over-union matching to produce a 3D label volume.

From labelled masks, NucleiSky extracts nuclear centroids and per-nucleus descriptors for use with feature-driven matchers. These descriptors include region-based shape measures and local neighbourhood context, such as k-nearest-neighbour distances and density estimates in micrometre space. For 3D labels, similar centroid and neighbourhood features are extracted using SimpleITK when available, or provided directly as centroid tables. In both cases, consistency with the provided voxel calibration is preserved.

### Graph matcher

The graph matcher constructs a k-nearest-neighbours graph from nuclear centroids measured in micrometres and generates a rotation-resistant node descriptor that encodes locally normalised neighbour distances and relative angles. This descriptor, derived from the graph, can be combined with per-nucleus morphological and neighbourhood feature vectors obtained from the segmentation mask. Candidate matches between query and reference nuclei are identified via nearest-neighbour search in the combined feature space. Optionally, multiple candidates per nucleus may be retained to handle ambiguous matches. These candidates can be filtered using ratio-based ambiguity tests and robust feature-distance thresholds. The matcher then proposes similarity transforms by sampling correspondence triplets in a RANSAC-like process, discarding degenerate triplets with very small triangle areas and evaluating hypotheses based on spatial inlier consensus. The best hypothesis can be refined using the full set of inliers and further adjusted via iterative closest-point-like updates to the similarity transform. To avoid overly strict matching in dense tissues, the graph matcher enforces a minimum inlier radius based on the typical nearest-neighbour spacing in the reference constellation.

### Triangle matcher

The triangle matcher derives local geometric descriptors from triangles within a k-nearest-neighbour neighbourhood. For each nucleus, triangles are formed by anchoring on a local neighbour and combining it with additional neighbours. Near-degenerate configurations below a predefined area threshold are discarded. Triangle descriptors are aggregated per nucleus and normalised using statistics from the reference constellation. Candidate correspondences are found via a nearest-neighbour search in triangle-descriptor space, and similarity transforms are proposed via RANSAC-style sampling of correspondence triplets. Candidate hypotheses are filtered by configured scale and rotation limits and scored by spatial consensus among inliers. The best hypothesis can optionally be refined with ICP-like updates.

### Quad matcher

The quad matcher analyses multiple local four-point configurations around each nucleus by selecting neighbour triplets from a k-nearest-neighbours set. Each configuration is encoded with a rotation- and scale-invariant descriptor derived from normalised distances and a deterministic neighbour ordering. Near-degenerate quads are discarded using a minimum-area criterion. Query quad descriptors are matched to reference quad descriptors via nearest-neighbour retrieval in descriptor space, yielding candidate configuration pairs. Candidate similarity transforms are derived from matched quads. Hypotheses from multiple three-point subsets of each quad may be generated to tolerate a single incorrect neighbour within a local configuration. These hypotheses are screened using scale and optional rotation constraints and scored based on inlier consensus. Early stopping can be applied once an inlier target is reached, and the final transform can be refined using the inliers.

### Geometric hashing matcher

The geometric hashing matcher accelerates localisation in large reference sets by indexing local coordinate frames in a hash table. In 2D, local frames are generated from short anchor pairs selected from k-nearest-neighbour lists. Additional neighbours are encoded in a scale-normalised polar representation relative to the anchor-defined frame. These signatures are quantised into radial and angular bins and stored in a hash map with per-bin limits to manage memory and runtime. During querying, similar local arrangements are sampled from the query set. Hash bins are searched using a configurable neighbour-bin radius to reduce quantisation sensitivity, and retrieved hash entries are ranked and tested to generate candidate transform hypotheses. These hypotheses are filtered by scale and optional rotation bounds and scored using spatial inlier consensus. An optional pre-test on a subset of nuclei allows early rejection of weak hypotheses. The selected transform can be refined through iterative closest-point–like updates.

### Adaptive matching controller

To accommodate variations in nuclear density, segmentation quality, and query size, NucleiSky provides an adaptive controller that runs a sequence of matchers and stops at the first successful localisation. The controller selects the order of matchers based on the number of query nuclei and skips matchers whose prerequisites are unmet. For example, the graph matcher is skipped when per-nucleus feature vectors are unavailable. If no matcher succeeds, the controller returns the highest-scoring candidate under a policy that prioritises transforms with high inlier fractions and low mean inlier errors, enabling best-effort localisation while preserving diagnostic information about failure modes.

### NucleiSky3D

NucleiSky extends constellation matching to volumetric datasets by representing nuclei as 3D centroids in micrometre space and estimating a 3D similarity transform comprising a uniform scale, a 3D rotation, and a translation. The 3D pipeline features two matchers, the pyramid matcher and 3D geometric hashing. The pyramid matcher constructs local tetrahedra from k-nearest-neighbour neighbourhoods and encodes each with a descriptor based on normalised edge-length patterns and a volume term normalised by the typical edge scale, providing invariance to rotation and uniform scaling while rejecting degenerate tetrahedra. Candidate tetrahedra are matched in descriptor space using mutual filtering, and 3D similarity transforms are estimated by sampling tetrahedron correspondences and scoring them via inlier consensus. The 3D geometric hashing matcher defines local 3D coordinate frames from three anchor points and encodes the relative coordinates of a fourth point within that frame, which are quantised into bins for hash-based retrieval. Candidate hypotheses are generated by querying the hash map with configurations, filtering by scale and optional rotation limits, and scoring them using inlier consensus, with optional pre-test pruning and iterative refinement. An adaptive 3D controller chooses between pyramid-first and 3D geometric hashing-first strategies based on the query constellation size, using a threshold of 1,000 nuclei in the current implementation.

### Outputs, export, and reproducibility

For each successful run, NucleiSky provides estimated similarity transform parameters and match-quality metrics, including inlier statistics and localisation error estimates derived from nearest-neighbour distances in micrometre space. The pipeline can export aligned overlays and derived outputs to support validation and downstream analysis. For example, predicted query footprints in reference pixel coordinates and transform records suitable for reuse across sessions. When segmentation masks are provided, they can be saved alongside transforms and overlays to enhance reproducibility and facilitate auditing of segmentation-dependent effects.

### Matcher benchmarking and performance analysis

We evaluated NucleiSky using a patch-localisation task in which square query fields of view were cropped from a larger reference image, ensuring known ground-truth coordinates for each query. For each reference image, landmark centroids were precomputed and stored in both pixel and micrometre units. Query landmarks were defined as those whose centroid coordinates fell within the crop. Their coordinates were then transformed into the query patch’s local coordinate system and converted to micrometres using the image’s pixel size, whereas reference landmarks remained in the coordinate system of the full reference image. Because the query and reference shared the same pixel calibration in this benchmark, correct localisation meant recovering the crop’s translation in the reference image.

For the localisation benchmark, square query patches of increasing sizes were tested. For each patch size, 50 candidate query locations were generated deterministically and reused across matchers. Patch origins were anchored to landmarks to ensure informative configurations: a landmark was sampled, a random position within the patch was selected, and the resulting top-left coordinate was clipped to the image bounds. Duplicate candidates were removed and, when enabled, candidates with fewer than 3 landmarks were discarded. The graph, quad, triangle, hashing, and adaptive matchers were evaluated on the same query patches. For each patch, we recorded the estimated translation, matcher-reported success, and match-quality statistics, including inlier fraction and mean inlier error. Translation accuracy was measured as the Euclidean distance between the estimated translation and the ground-truth crop origin, in pixels. When a translation was available, we also computed the structural similarity index measure (SSIM) between the query patch and the predicted reference crop after identical percentile-based intensity normalisation. Localisation was considered successful on the standard benchmark only when the matcher reported success and all predefined quality thresholds were met: inlier fraction 0.80, SSIM 0.80, and translation error <20 pixels. Success probabilities were summarised as a function of query patch size and as a function of the number of landmarks in the query field of view. Confidence intervals were calculated using the Wilson score interval. For nuclei-count summaries, observed success probabilities were binned by landmark count and plotted on a logarithmic x-axis; logistic fits in log10 (landmark count) space were used to estimate the number of landmarks required to reach 90% localisation probability.

To evaluate sensitivity to segmentation errors, we conducted a separate benchmark using 512-pixel query patches. Candidate patches were selected to contain at least 50 clean landmarks before perturbation. Adaptive mode was excluded from this benchmark because the goal was to compare the robustness of individual matchers under controlled landmark corruption. In addition, the adaptive controller uses internal success criteria that are not appropriate when false-positive or false-negative landmarks are deliberately introduced. We therefore evaluated the quad, triangle, and hashing matchers individually. Two segmentation-error modes were tested independently. To mimic missed detections, a specified fraction of true landmarks was randomly removed from the set of centroids. To mimic spurious detections, synthetic centroids were added uniformly within the relevant image domain. Extra detections were expressed relative to the number of true detections after any removal; for example, 100% extra detections added one synthetic centroid per true centroid, whereas 1000% added ten synthetic centroids per true centroid. Missed-detection levels of 0, 10, 20, 40, 60, 80, 90, and 95% were tested, as were extra-detection levels of 0, 10, 25, 50, 100, 200, 500, and 1000%.

Segmentation perturbations were applied to both the query and reference centroid sets. For query perturbations, synthetic detections were sampled uniformly within the 512-pixel query patch. For reference perturbations, synthetic detections were uniformly sampled across the full reference image, with an upper limit on the number of added reference points to avoid excessive memory usage and runtime. Patches or perturbed centroid sets with fewer than three landmarks after perturbation were recorded as failures. For the segmentation-error benchmark, matcher-reported success and inlier fraction were recorded only as diagnostics and were not used to define benchmark success. This was necessary because removing true landmarks changes the maximum attainable inlier fraction, while adding spurious detections changes the consensus denominator and can distort internal success flags. Instead, success was defined solely by image-level agreement: the predicted reference crop was compared with the original unperturbed query image using SSIM after identical percentile-based intensity normalisation, and a trial was considered successful when SSIM 0.80. Success probabilities were reported with 95% Wilson confidence intervals. Extra-detection levels were plotted as tested categories because the perturbation series was intentionally uneven.

Runtime was measured as both matcher time and total per-patch time, with total time including patch extraction and metric calculation. Benchmarks were run using the NucleiSkyApp benchmarking notebooks on a laptop workstation running Ubuntu 26.04 LTS with an Intel Core i7-13700H CPU, 64 GB RAM, and an NVIDIA GeForce RTX 4060 Laptop GPU.

### 3D benchmark and performance analysis

We evaluated NucleiSky3D using a volumetric patch-localisation task similar to the 2D benchmark. Query subvolumes were cropped from a larger reference volume, providing known ground-truth positions for each query. For each reference, 3D landmark centroids were extracted from label masks and stored in both pixel and calibrated micrometre coordinates. Query landmarks were defined as nuclei whose centroids fell within the cropped subvolume. Their coordinates were transformed into the query’s local coordinate system by subtracting the crop origin and converting to micrometres using the voxel size, whereas the reference landmarks remained in the coordinate system of the full reference volume. Because the query and reference came from the same calibrated volume in this benchmark, correct localisation corresponded to recovering the 3D crop translation within the reference.

For each tested subvolume size, candidate query locations were generated deterministically and reused across matchers. Patch origins were anchored to landmarks to ensure informative configurations: a landmark was sampled, assigned a random position within the subvolume, and the resulting Z, Y, X crop origin was clipped to the reference bounds. Duplicate candidates were removed, and subvolumes with fewer than three landmarks were discarded before matching. Patch sizes were specified in Z, Y, X order and automatically filtered to evaluate only subvolumes that fit within the loaded reference volume. Unless otherwise stated, 100 candidate subvolumes were tested per patch size.

We evaluated the pyramid, 3D geometric hashing, and adaptive 3D matchers on the same query subvolumes. The adaptive controller used the pyramid and 3D geometric hashing matchers, stopping once successful localisation was achieved. For each query, we recorded the matcher-reported success, the estimated 3D translation, the inlier fraction, the mean inlier error, the estimated scale, the estimated rotation magnitude, the per-axis translation errors, and the runtime. Translation accuracy was measured by comparing the estimated translation with the known crop origin in both calibrated coordinates and pixels. Total runtime included patch extraction, matching, and metric calculation, whereas matcher runtime measured the matching call itself.

For plotting and summary statistics, successful 3D localisation was defined as matcher-reported success, an inlier fraction ≥ 0.50, and a 3D translation error <20 pixels. The XY maximum-intensity-projection SSIM between the query subvolume and the predicted reference subvolume was computed after identical percentile-based intensity normalisation. It was recorded only as an image-level diagnostic and not used as a volumetric success criterion. Success probabilities were summarised as a function of query subvolume size and of the number of landmarks in the query subvolume. Confidence intervals were calculated using the Wilson score interval. Landmark-count summaries were plotted on a logarithmic x-axis, and logistic fits in log10(landmark count) space were used to estimate the number of landmarks required to reach 90% localization probability. Benchmark results were check-pointed separately for each matcher and patch size, allowing interrupted runs to resume from existing result files.

Runtime was measured as both matcher time and total per-patch time, with total time including patch extraction and metric calculation. Benchmarks were run using the NucleiSkyApp benchmarking notebooks on a laptop workstation running Ubuntu 26.04 LTS with an Intel Core i7-13700H CPU, 64 GB RAM, and an NVIDIA GeForce RTX 4060 Laptop GPU.

### Cells

MCF10 DCIS.com cell lines (DCIS.com; invasive T24 c-Ha-ras oncogene-transfected breast epithelial cells (Miller et al., 2000)) were cultured in a 1:1 mix of DMEM (Merck) and F12 (Merck) supplemented with 5% horse serum (GIBCO BRL, Cat Number: 16050122), 20 ng/ml human EGF (Merck, Cat Number: E9644), 0.5 µg/ml hydrocortisone (Merck, Cat Number: H0888-1G), 100 ng/ml cholera toxin (Merck, Cat Number: C8052-1MG), 10 µg/ml insulin (Merck, Cat Number: I9278-5ML), and 1% (vol/vol) penicillin/streptomycin (Merck, Cat Number: P0781-100ML) at 37°C, 5% CO_2_. DCIS.com cells were provided by J.F. Marshall (Barts Cancer Institute, Queen Mary University of London, London, England, UK).

Human Umbilical Vein Endothelial Cells (HUVECs) (PromoCell, C-12203) were cultured in ready-to-use Endothelial Cell Growth Medium (ECGM) (PromoCell, C-22010 and C-39215), supplemented with 1% penicillin-streptomycin (Sigma-Aldrich, P0781). Human pulmonary microvascular endothelial cells (HPMEC; PromoCell, C-12281) were cultured in ready-to-use Endothelial Cell Growth Medium MV (ECGM MV; PromoCell, C-22020 and C-39225) supplemented with 1% penicillin-streptomycin (Sigma-Aldrich, P0781). Primary endothelial cells from “P0” (commercial vial) were expanded to a P3 stock and stored at −80°C until used to standardise experimental replicates. Vials were thawed, cells were seeded into a 10-cm culture dish, and the dish was incubated for at least 2 days.

Pancreatic ductal adenocarcinoma SU.86.86 cells (ATCC CRL-1837) and AsPC-1 cells (ATCC CRL-1682) were cultured in RPMI 1640 supplemented with 10% fetal bovine serum (FBS; BioWest, S1810), 1% L-glutamine (Sigma-Aldrich, G7513) and 1% penicillin-streptomycin (Sigma-Aldrich, P0781). AsPC-1 and SU.86.86 cells were authenticated by the Leibniz Institute DSMZ (Deutsche Sammlung von Mikroorganismen und Zellkulturen) via STR profiling. All cell lines were routinely tested for mycoplasma infections and found to be free of mycoplasma.

### Dataset 1: tiled widefield image of a DAPI-stained HUVEC monolayer

HUVECs were seeded in an ibidi chamber to form a confluent endothelial monolayer, then fixed with 4% paraformaldehyde (ThermoFisher Scientific, 28908) and stained with DAPI (ThermoFisher Scientific, D1306) to label nuclei. Widefield fluorescence images were captured on a Nikon Eclipse Ti2-E inverted microscope, controlled by Nikon NIS-Elements AR (v6.10), using a 10×/0.3 Plan Fluor objective and a Hamamatsu Orca Flash4.0 V3 sCMOS camera. Illumination was provided by a Lumencor Spectra/AuraII (Spectra X) light engine. The dataset used for benchmarking was acquired as a stitched mosaic in NIS-Elements and exported for analysis. For the nuclei channel, excitation was delivered via the 395 nm LED line, and emission was collected through the Nikon DAPI filter set. The camera was operated with 2×2 binning and 12-bit acquisition, with data stored in a 16-bit container. The reference image used for benchmarking measured 1,266 × 2,970 pixels, corresponding to 1,671.8 µm × 3,922.0 µm with a pixel size of 1.3205 µm per pixel. For benchmarking, nuclei were segmented using intensity thresholding (implemented via NucleiSky’s threshold-and-watershed backend), identifying 4,387 nuclei. This dataset has been deposited on Zenodo Hidalgo Cenalmor et al. (2026).

### Dataset 2: spinning-disk montage of an HUVEC monolayer co-incubated with SU.86.86 cells

HUVECs were seeded in an ibidi chamber to form a confluent endothelial monolayer, after which SU.86.86 cells were added for 2 h. Monolayers were washed, fixed with 4% paraformaldehyde (ThermoFisher Scientific, 28908), and stained with DAPI (ThermoFisher Scientific, D1306) to label nuclei. Imaging was performed on a 3i / Zeiss Marianas CSU-W1 spinning-disk confocal inverted microscope equipped with a 20×/0.8 Plan-Apochromat air objective and an Orca Flash4.0 sCMOS camera, controlled by 3i SlideBook (v6). Acquisition was performed using the 405-nm channel for nuclei. The exposure time for the 405-nm channel was 50 ms. Z-stacks were acquired and stitched into a montage, and the dataset was analysed as a maximum-intensity projection. The resulting projected montage had dimensions of 5,520 × 5,520 pixels with a calibrated field of view of 3,500.488 µm × 3,500.488 µm (pixel size 0.6341 µm/pixel). For benchmarking, nuclei were segmented from the DAPI channel using the Cellpose backend (Cellpose-SAM configuration), identifying 14,418 nuclei. This dataset has been deposited on Zenodo Hidalgo Cenalmor et al. (2026).

### Dataset 3: public H&E whole-slide image of mouse colon

Dataset 3 is an H&E bright-field image of a mouse colon section, obtained from the 10x Genomics datasets portal (dataset: “Fresh Frozen Mouse Colon with Xenium Multimodal Cell Segmentation”; https://www.10xgenomics.com/datasets). The image was provided as an OME-TIFF RGB image (8-bit; three channels corresponding to RGB) with dimensions of 32,201 × 28,329 pixels and a calibrated sampling of 0.27377 µm/pixel (corresponding to an imaged area of 8.82 mm × 7.76 mm). For benchmarking, nuclei were segmented from the H&E image using the InstanSeg backend, identifying 247,107 nuclei.

### Dataset 4: widefield tiled imaging of fixed Escherichia coli cells

Escherichia coli cells were cultured under standard conditions at 37°C in LB broth, sedimented onto glass coverslips, fixed with 4% paraformaldehyde for 10 min, and stained with DAPI (ThermoFisher Scientific, D1306) to label bacterial nucleoids. Imaging was performed on a Zeiss Axio Observer 7 inverted microscope equipped with a dual-filter wheel and a Prime 95B camera, using a Plan-Apochromat 40×/0.95 Korr M27 objective with a 1.6× optovar, resulting in an effective magnification of 64× and a calibrated pixel size of 0.171875 µm/pixel. For benchmarking, the stitched reference image used measured 11,788 × 11,788 pixels, corresponding to approximately 2.03 × 2.03 mm. DAPI-stained bacterial nucleoids were segmented using the Cellpose-SAM backend in NucleiSky, identifying 81,379 detected objects. This dataset has been deposited on Zenodo Hidalgo Cenalmor et al. (2026).

### Datasets 5 and 6: DCIS.com monolayers for cross-scale localisation

Datasets 5 and 6 consist of confluent monolayers of DCIS.com cells that transiently express MYO10^HMM^-GFP (a truncated Myosin-X construct used to label filopodia tips (Grobe et al., 2026)) and are stained to visualise the plasma membrane and nuclei. Briefly, DCIS.com cells transiently expressing MYO10^HMM^-GFP were seeded into ibidi 2-well culture inserts (ibidi, 81176) positioned at the centre of MatTek 35-mm gridded glass-bottom dishes (P35G-1.5-14-CGRD; MatTek) coated with poly-L-lysine and fibronectin. Cells were fixed for 25 min at room temperature in 2% formaldehyde and 0.05% glutaraldehyde prepared in 0.1 M HEPES supplemented with 2 mM CaCl and 2 mM MgCl. Samples were stained with wheat germ agglutinin (WGA; 1:1000) to label the cell surface and with DAPI (ThermoFisher Scientific, D1306) to label nuclei. Between steps, samples were washed three times with 0.1 M HEPES supplemented with 2 mM CaCl and 2 mM MgCl. Images were acquired on a Zeiss LSM 880 inverted microscope equipped with an Airyscan detector and controlled with Zeiss ZEN 2.3. Airyscan processing was performed in standard super-resolution mode, and tiled acquisitions were stitched in ZEN as indicated below.

Dataset 5 includes a 20× overview mosaic and a 63× query. For the reference overview, a tiled acquisition was performed with a Plan-Apochromat 20×/0.8 air objective and processed using Airyscan/Zen stitching and fusion, yielding a field of view of 6,966 × 7,143 pixels and a calibrated pixel size of 0.179214 µm/pixel (X, Y). For the high-magnification query, a confocal Z-stack was captured with a C Plan-Apochromat 63×/1.4 oil objective, yielding 2,372 × 2,372 pixels at 0.056893 µm/pixel (X, Y) and a Z sampling of 0.184964 µm. For localisation, the stack was converted to a 2D view using a maximum-intensity projection of the nuclei channel.

Dataset 6 includes a 10× overview mosaic and a 63× query. The reference overview was captured with a Plan-Apochromat 10×/0.3 air objective and processed using Airyscan/Zen stitching and fusion. This produced a fused mosaic of 6,844 × 6,904 pixels, with a calibrated pixel size of 0.2679 µm/pixel, covering approximately 1.83 × 1.85 mm. The high-magnification query involved acquiring a 63× Z-stack with a C Plan-Apochromat 63×/1.4 oil objective, yielding an image of 2,372 × 2,372 pixels with voxel sampling of 0.0316 × 0.0316 × 0.1850 µm^3^ (X, Y, Z) across 95 Z-planes. For localisation, a single Z-plane from this stack was selected.

For both datasets, nuclei were segmented independently in the reference and query images using the InstanSeg backend. To harmonise nuclear appearance across magnifications, images were rescaled with the NucleiSky rescaling option before segmentation.

Both datasets have been deposited on Zenodo Hidalgo Cenalmor et al. (2026).

### Dataset 7: live-to-fixed imaging using bright-field-derived nuclear labels

Dataset 7 was used to test live-to-fixed registration between a label-free bright-field movie and a post-fixation fluorescence tile scan. The experiment was conducted using FlowVision, in which circulating pancreatic cancer cells are perfused over an endothelial monolayer under controlled flow conditions and imaged using fast transmitted-light bright-field microscopy (Follain et al., 2026). For live imaging, AsPC-1 cells in suspension (2 × 105 cells/ml) were perfused over a confluent HPMEC monolayer within a fibronectin-coated microfluidic channel, following the FlowVision endothelial arrest assay, in which circulating cells are exposed to controlled capillary-like flow over an endothelial monolayer. The assay uses a microfluidic channel connected to a perfusion system to deliver cancer cells over endothelial monolayers under defined flow conditions while imaging in a temperature- and CO-controlled microscope chamber (Follain et al., 2026). Live imaging was performed in bright-field mode to avoid fluorescence phototoxicity and preserve downstream endpoint staining options. In this experiment, cells were perfused for 8 min, with the flow speed changed every 2 min from 300 µL/min to 200 µL/min, then to 100 µL/min, followed by a return to 300 µL/min as a wash step to assess adhesion stability. The live bright-field movie used for NucleiSky registration was acquired as a 16-bit time series of 12,000 frames, with a 40 ms frame interval. Each frame measured 2,048 × 2,044 pixels, corresponding to 664.6 × 663.3 µm, with a calibrated pixel size of 0.3245 µm/pixel. Because nuclei were not directly acquired as a fluorescent channel during live imaging, synthetic nuclear labels were generated from the bright-field data using the pix2pix artificial-labelling model developed for FlowVision (Follain et al., 2026; Isola et al., 2016). This model was trained on paired bright-field and fluorescence images to infer DAPI-like nuclear signal from bright-field input images. The pix2pix-derived nuclear image used for NucleiSky analysis measured 1,024 × 1,024 pixels and served as the query image. It was segmented using the Cellpose-SAM backend in NucleiSky to generate the live-derived nuclear centroid constellation.

After live imaging, the same sample was immediately fixed for 10 min with 4% PFA at a flow rate of 200 µL/min. The PFA was washed with PBS for 10 min at a flow rate of 200 µL/min, then stored overnight at 4 °C. Immunostaining for TOMM20 was performed prior to Cell Painting staining. Cells were permeabilised with 0.2% Triton X-100 for 10 min at room temperature on a rocker, washed three times with 150 µL PBS, and quenched with 0.2 M glycine for 30 min at room temperature on a rocker. After three washes with PBS, cells were incubated with the mouse anti-TOMM20 antibody (1:200; Abcam, cat. no. ab56783) for 1 h at room temperature on a rocker. Cells were then washed three times with PBS and incubated with Alexa Fluor 647 goat anti-mouse secondary antibody (1:400; Thermo Fisher Scientific, cat. no. A21235) for 30 min at room temperature on a rocker. After three further PBS washes, cells were stained with the Image-iT Cell Painting Kit (Invitrogen, cat. No. I65000) according to the manufacturer’s protocol, omitting MitoTracker. Briefly, cells were incubated with Cell Painting blocking solution (1% BSA + 0.1% Triton X-100 in HBSS, filtered through a 0.2 µm membrane) for 20 min at room temperature on a rocker, washed three times with PBS, and incubated for 30 min at room temperature on a rocker with the Cell Painting dye mixture containing Hoechst 34580, Concanavalin A–Alexa Fluor 488, Wheat Germ Agglutinin–Alexa Fluor 555, Alexa Fluor 568 phalloidin, and SYTO 14. The final working concentrations were 5 µg/mL Hoechst 34580, 100 µg/mL Concanavalin A–Alexa Fluor 488, 3 µg/mL Wheat Germ Agglutinin–Alexa Fluor 555, 50 ng/mL Alexa Fluor 568 phalloidin, and 6.5 µM SYTO 14. Cells were washed three times with PBS and stored at 4°C, protected from light, until imaging.

Images were acquired on a 3i/Zeiss Marianas CSU-W1 spinning-disk confocal microscope equipped with a Yokogawa CSU-W1 scanner, a Hamamatsu ORCA-Flash4.0 sCMOS camera, and a Zeiss Plan-Apochromat 20×/0.8 air objective. Images were acquired using Slidebook 6 software with 1×1 binning, a calibrated pixel size of 0.317 µm/pixel, and a Z-step size of 0.630 µm. Each field of view measured 649 × 649 µm (2048 × 2048 pixels). The four-channel Z-stacks were acquired as a 5 × 5 tile montage comprising 25 fields of view, and the montage was generated with Slidebook 6. The fixed nuclei (Hoechst 34580) reference was exported as a maximum-intensity projection and used as the reference image for NucleiSky registration. The reference image measured 9,421 × 9,421 pixels, corresponding to 2,985.3 × 2,985.3 µm, with a calibrated pixel size of 0.3169 µm/pixel. The image was 16-bit and segmented from the nuclei channel using the Cellpose-SAM backend in NucleiSky.

For NucleiSky analysis, the pix2pix-derived live nuclear centroids served as the query constellation, and the Hoechst-derived fixed nuclear centroids as the reference constellation. Both centroid sets were converted to calibrated physical coordinates before matching. NucleiSky was then used to estimate a 2D similarity transform that mapped the live bright-field-derived query field of view onto the fixed fluorescence reference tile scan. The resulting transform located the live-imaged field of view within the post-fixation reference image and enabled transfer of the alignment to additional endpoint fluorescence channels.

This dataset has been deposited on Zenodo Hidalgo Cenalmor et al. (2026).

### Dataset 8: dataset used for smart microscopy

Dataset 8 was used to test real-time smart microscopy re-targeting across microscopes. The dataset consisted of three acquisition stages: a query image acquired on Microscope A, low-magnification overview images acquired during automated scanning on Microscope B, and high-resolution images acquired after NucleiSky identified the matched field of view. HUVEC cells were seeded on 13 mm circular coverslips coated with fibronectin (10 µg/mL) at a density of 50,000 cells per coverslip and cultured in 12-well plates. After 48 h, cells were fixed with 4% PFA in culture medium for 10 min at room temperature, washed three times with PBS, and stained using the Image-iT Cell Painting Kit, omitting MitoTracker. Cells were incubated with Cell Painting blocking solution for 20 min at room temperature on a rocker, washed three times with PBS, and incubated for 30 min at room temperature with Hoechst 34580, Concanavalin A–Alexa Fluor 488, Wheat Germ Agglutinin–Alexa Fluor 555, Alexa Fluor 568 phalloidin, and SYTO 14 at final working concentrations of 5 µg/mL, 100 µg/mL, 3 µg/mL, 50 ng/mL, and 6.5 µM, respectively. Cells were then washed three times with PBS and stored at 4°C protected from light until imaging. The query image was acquired on a Nikon Eclipse Ti2-E inverted microscope using NIS-Elements AR (v6.10) and a 20×/0.75 Plan Apo Lambda objective, with a Hamamatsu Orca Flash4.0 V3 camera. Illumination was provided by a Lumencor Spectra/AuraII light engine. Images were acquired in three fluorescence channels: Hoechst342, AF568, and SYTO 14, using excitation wavelengths of 395, 550, and 475 nm, respectively. All channels were acquired with a 100 ms exposure time. The query was acquired as a 7-plane Z-stack with 0.9 µm spacing. Each plane measured 2,044 × 2,048 pixels, corresponding to 663.28 × 664.58 µm, with a calibrated pixel size of 0.3245 µm/pixel. For NucleiSky matching, the nuclei channel was converted to a maximum-intensity projection and used as the query image.

Overview scanning and matched high-resolution imaging were performed on a Nikon Ti2-E Crest V3 inverted microscope controlled by NIS-Elements AR (v6.20). Low-magnification overview images were acquired using a 4×/0.2 CFI Plan Apochromat Lambda D objective and a Photometrics Kinetix camera. Overview images were acquired from the nuclei channel. Each overview position was captured as a 5-plane Z-stack with 12.5 µm spacing. Within the NIS-Elements JOBS workflow, each overview stack was processed using a maximum-intensity-projection block followed by a Denoise.ai block before being passed to NucleiSky. The processed overview images measured 2,048 × 2,048 pixels, corresponding to 3,430.51 × 3,430.51 µm, with a calibrated pixel size of 1.6751 µm/pixel. NucleiSky compared the processed overview image with the query image to identify the matching nuclear constellation. When no acceptable match was detected, the JOBS workflow advanced to the next overview position. When a match was detected, the microscope moved to the predicted position and, using a 20×/0.8 CFI Plan Apochromat Lambda D objective and a Photometrics Kinetix sCMOS camera, acquired a three-channel image (Hoechst342, AF568, and SYTO 14, excited at 406, 545, and 476 nm, respectively) as a 3 × 3 stitched mosaic with 30% overlap and a 9-plane Z-stack with 0.8 µm spacing. This resulted in a 5,735 × 5,735-pixel image with a calibrated pixel size of 0.3332 µm/pixel.

This dataset has been deposited on Zenodo Hidalgo Cenalmor et al. (2026)

### Dataset 9: Platynereis-Nuclei-CBG

Dataset 9 was the publicly available Platynereis-Nuclei-CBG dataset (Lalit et al., 2022), used to evaluate 3D localisation with pre-existing nuclear instance masks. The reference volume measured 700 × 660 × 113 pixels (X, Y, Z) and contained 392 segmented nuclei after conversion of the 3D label mask into nuclear centroids. Pre-existing 3D instance masks were used directly; no additional nuclear segmentation was performed. Label objects were converted to calibrated 3D centroids and used as the reference landmark set for NucleiSky3D benchmarking.

### Dataset 10: DAPI-stained mouse brain 3D benchmark dataset

Dataset 10 was prepared and imaged as previously reported (Ball et al., 2024). Briefly, brains from female mice (Hsd: Athymic Nude-Foxn/1nu, Envigo) were dissected, rinsed in PBS, and fixed overnight at 4°C in 4% paraformaldehyde (ThermoFisher Scientific, 28908). Fixed brains were embedded in low-melting-point agarose, and 100–200 µm sections were cut using a vibratome. Brain slices were permeabilised, blocked and stained. Samples were mounted in ProLong Glass Antifade Mountant (P36980; Thermo Fisher Scientific). Images were acquired on a Marianas spinning-disk confocal microscope using a Yokogawa CSU-W1 spinning-disk unit on an inverted Zeiss Axio Observer Z1 microscope controlled by SlideBook 6, with an Orca Flash 4 sCMOS camera and a 63×/1.4 oil Plan-Apochromat objective. The analysed volume measured 1,840 × 1,840 × 224 pixels (X, Y, Z), corresponding to 185.2 × 185.2 × 67.2 µm, with voxel sampling of 0.1007 × 0.1007 × 0.2998 µm (X, Y, Z). The DAPI channel was used to generate the 3D nuclear landmark reference. Individual Z-planes were segmented using the Cellpose-SAM backend in NucleiSky, and 3D nuclear objects were reconstructed by merging labels across adjacent Z-slices using the NucleiSky 2.5D label-linking workflow. This produced 864 segmented nuclei, which were reduced to calibrated 3D centroids for benchmarking. Query subvolumes of increasing size were cropped from this reference and re-localised using the pyramid, 3D geometric hashing and adaptive NucleiSky3D matchers. This dataset has been deposited on Zenodo Hidalgo Cenalmor et al. (2026).

## Data availability

All datasets used in this study are either already publicly available (as indicated) or deposited on Zenodo Hidalgo Cenalmor et al. (2026). The authors declare that the data supporting the findings of this study are available within the article and from the authors upon request. Any additional information needed to reanalyse the data reported in this paper is available from the corresponding authors.

## Code availability

All code used in this study is available under the MIT license at https://github.com/CellMigrationLab/NucleiSky.

## Manuscript preparation

Figures were prepared using Fiji (Schindelin et al., 2012) and Inkscape. PlotsOfData was used to generate the box plot shown in Fig. S1 (Postma and Goedhart, 2019). GPT-5.5 (OpenAI) and Grammarly (Grammarly, Inc.) served as writing aids during manuscript preparation. The author also edited and validated all sections of the text. The PDF version of this manuscript was formatted with Rxiv-Maker (Saraiva et al., 2025b).

## Author contributions

**Conceptualisation**: I.H.C, J.K.A, P.N., G.J. **Methodology**: I.H.C, A.O.O., G.J. **Reagents**: G.J. **Software**: G.J. and I.H.C **Formal Analysis**: I.H.C, A.O.O., G.J. **Investigation**: I.H.C, A.O.O, C.P., G.J. **Writing - Original Draft**: G.J. **Writing - Review and Editing**: Everyone. **Visualisation**: G.J., I.H.C. **Supervision**: G.J., R.H., M.D.R **Funding Acquisition**: G.J.

## Acknowledgments

This study was funded by the Research Council of Finland (338537, 371287, and 374180 to G.J.), the Sigrid Juselius Foundation (to G.J.), the Cancer Society of Finland (Syöpäjärjestöt; to G.J.), and the Solutions for Health strategic funding for Åbo Akademi University (to G.J.). The research was also supported by the InFLAMES Flagships Program of the Research Council of Finland (decision numbers: 337530, 337531, and 357910). G.J. is supported by the Finnish Cancer Institute (K. Albin Johansson Professorship). I.H.C. is funded by the Finnish Doctoral Program Network in Artificial Intelligence, AI-DOC (decision number VN/3137/2024-OKM-6). P.N. is supported by the Fru Berta Kamprads Foundation (FBKS-2024-8 - (584). R.H., M.D.R., and C.P. acknowledge funding from the European Research Council (ERC) under the European Union’s Horizon 2020 research and innovation programme (SelfDriving4DSR, grant agreement No. 101001332). R.H. is further supported by the European Union through Horizon Europe (RT-SuperES, grant agreement No. 101099654); by a European Molecular Biology Organization (EMBO) Installation Grant (EMBO-2020-IG-4734); by a Chan Zuckerberg Initiative Visual Proteomics Imaging award (vpi-0000000044); by a joint Wellcome, Chan Zuckerberg Initiative, and Kavli Foundation Essential Open Source Software for Science Cycle 6 award (Wellcome 313383/Z/24/Z; CZI EOSS6-0000000260); and by the La Caixa Foundation (CaixaResearch Health 2025, VirusAwareScopes, HR25-00453). R.H. and C.P. were also supported by Fundação para a Ciência e a Tecnologia, through the MOSTMICRO-ITQB RD Unit (UID/PRR/4612/2025) and the LS4FUTURE Associated Laboratory. The Chan Zuckerberg Initiative awards are made through the Chan Zuckerberg Initiative DAF, an advised fund of Silicon Valley Community Foundation. This study was funded by the European Union; however, the views and opinions expressed are those of the authors only and do not necessarily reflect those of the European Union or the granting authority. Neither the European Union nor the granting authority can be held responsible for them.

Imaging was performed at the Advanced Imaging Core of the Turku Bioscience Centre. This facility received support from Turku BioImaging, Biocenter Finland, and the Finnish Advanced Microscopy Node of Euro-BioImaging Finland (funded by the Research Council of Finland, FIRI 2023 grant decision numbers 359073 and 358879, and FIRI 2024 grant decision numbers 367582 and 367577).

## ABOUT THIS MANUSCRIPT

This work is licensed under CC BY 4.0.

## Supplementary Information

***Video 1. Smart microscopy relocalisation workflow in NIS-Elements AR***. Screen recording of the smart microscopy relocalisation workflow implemented using JOBS in NIS-Elements AR software, shown alongside (Fig. 5) to highlight the corresponding workflow step. The workflow aims to locate a query field-of-view image acquired on a different microscope than the microscope shown in the video. The video covers the complete process, including manual configuration of the acquisition area and input parameters, automated slide scanning, execution of NucleiSky to compare the query and reference images, and automated final image acquisition.

**Sup. Fig. S1.**
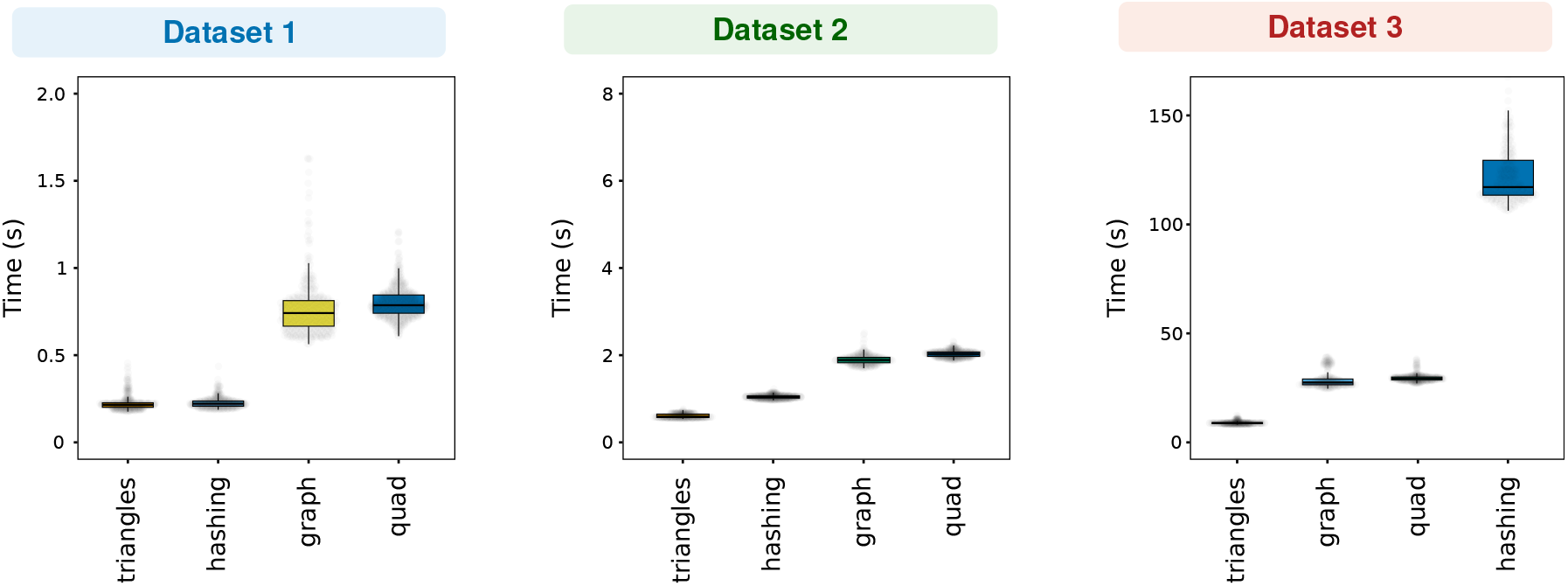
Benchmarking NucleiSky speed. Box plots show the distribution of execution times for each matcher to achieve a valid match across Datasets 1-3. To focus on true-positive localisation time, failed matching attempts are excluded from this analysis. In each dataset panel, the algorithms are ordered from left to right by increasing median execution time (fastest to slowest).

